# Coevolutionary dynamics of genetic traits and their long-term extended effects under non-random interactions

**DOI:** 10.1101/2020.03.10.985671

**Authors:** Charles Mullon, Joe Yuichiro Wakano, Hisashi Ohtsuki

## Abstract

Organisms continuously modify their living conditions via extended genetic effects on their environment, microbiome, and in some species culture. These effects can impact the fitness of current but also future conspecifics due to non-genetic transmission via ecological or cultural inheritance. In this case, selection on a gene with extended effects depends on the degree to which current and future genetic relatives are exposed to modified conditions. Here, we detail the selection gradient on a quantitative trait with extended effects in a patch-structured population, when gene flow between patches is limited and ecological inheritance within patches can be biased towards offspring. Such a situation is relevant to understand evolutionary driven changes in individual condition that can be preferentially transmitted from parent to offspring, such as cellular state, micro-environments (e.g., nests), pathogens, microbiome, or culture. Our analysis quantifies how the interaction between limited gene flow and biased ecological inheritance influences the joint evolutionary dynamics of traits together with the conditions they modify, helping understand adaptation via non-genetic modifications. As an illustration, we apply our analysis to a gene-culture coevolution scenario in which genetically-determined learning strategies coevolve with adaptive knowledge. In particular, we show that when social learning is synergistic, selection can favour strategies that generate remarkable levels of knowledge under intermediate levels of both vertical cultural transmission and limited dispersal. More broadly, our theory yields insights into the interplay between genetic and non-genetic inheritance, with implications for how organisms evolve to transform their environments.

## 1 Introduction

Genes often exert effects that extend beyond the organism in which they are expressed, for instance by modifying the physical environment (as with the building of nests or burrows), by altering ecological interactions (as when immunity genes influence an organism’s pathogens or microbiotic symbionts), or by creating cultural knowledge (as with the collection and dissemination of information about the environment; Dawkins, 1982; Lewontin, 1983; Odling-Smee et al., 2003; Bailey, 2012; Govaert et al., 2019). When genetic variation causes variation in some external characteristics that in turn leads to variation in reproductive success, these external characteristics can be considered as part of an organism’s extended phenotype (Dawkins, 1982). This opens a feedback where changes in the genetic composition of a population depend on external conditions themselves influenced by genes, so that adaptation involves changes not only in genetic characters, but also beyond the organisms that express these characters.

While feedbacks between genes and the environment can impact evolutionary dynamics in various ways (Robertson, 1991), their relevance for adaptation depends on the associations between genes, their extended effects and fitness (Dawkins, 1982, 2004; Brodie, 2005; Govaert et al., 2019). To see this, consider for instance a genetic locus that influences the quality of individual nests. For selection at this locus to causally depend on feedback effects, genetic variation must be linked to nest variation such that over generations, genes associated with “good” nests replicate at the expense of competitor genes associated with “bad” nests (to paraphrase Dawkins, 2004, p. 379). With this in mind, one consideration that is particularly relevant is whether the extended effects of genes extend further across generations, i.e., whether individuals can transmit elements of their extended phenotype to downstream generations via non-genetic pathways. This can occur under a variety of scenarios: material resources that have been modified by organisms are transferred to future generations; altered microbiomes are transmitted to offspring via physical contact; and cumulated cultural knowledge is passed down from older to younger individuals by imitation (Odling-Smee et al., 2003; Bonduriansky, 2012). In such cases, genes expressed in current individuals affect future generations through trans-generational extended effects, a phenomenon sometimes coined as “ecological inheritance” (Odling-Smee et al., 2003, or “cultural inheritance” when extended effects are specifically on cultural characteristics, Boyd and Richerson, 1985).

Selection on a gene with extended effects that can be ecologically transmitted depends on the degree to which current and future genetic relatives are exposed to conditions modified by a carrier of this gene (Lehmann, 2007, 2008). More specifically, the selection gradient on a quantitative character with inter-temporal effects can be expressed as the infinite sum of the marginal effects of a character change in one focal individual on the fitness of all current and future individuals in the population, each weighted by the genetic relatedness between the focal and the individual whose fitness is affected (e.g., eq. 2 in Lehmann, 2007). This kin selection perspective not only gives formal support to the notion that the adaptive significance of feedbacks between genetic traits and their extended effects is contingent on their association (Dawkins, 1982, 2004; Brodie, 2005), it also reveals that these associations depend on the genetic relatedness between individuals separated by multiple generations. But due to it generality, this selection gradient remains opaque about how different genetic and ecological processes influence the joint evolutionary dynamics of traits together with the conditions they modify (specifically, this requires characterising how different processes affect time-dependent relatedness and fitness effects).

In particular, it remains unclear how evolutionary dynamics are affected by the combined effects of limited gene flow between subpopulations and the mode of ecological transmission within subpopulations. Yet these two factors are expected to interact with one another in a way that is relevant for the feedback between genes and their extended effects. Indeed, if vertical transmission from parent to offspring (for e.g. due to maternal effects, Kirkpatrick and Lande, 1989; Mousseau and Fox, 1998; inheritance of acquired traits, Pál and Miklós, 1999; or preferential learning from parents, Boyd and Richerson, 1985) bolsters the association between genes and their extended effects (Day and Bonduriansky, 2011), this association is sapped by oblique transmission from non-parental individuals of the older generation to offspring (for e.g. owing to contagion of microbes, Brandvain et al., 2011; or oblique cultural learning, Boyd and Richerson, 1985). But where recipients of extended genetic effects via oblique ecological inheritance turn out to be genetic relatives due to limited gene flow, the feedback between genes and their extended effects can nevertheless materialise albeit indirectly through non-vertical kin (Lehmann, 2008). While a considerable variety of models has studied how feedbacks between genes and extended effects impacts evolutionary change, most are concerned with panmictic or well-mixed populations in the absence of any transmission bias (e.g., Bailey, 2012; Odling-Smee et al., 2013; Govaert et al., 2019, for reviews). Otherwise, evolutionary dynamics have been examined either under vertical transmission in panmictic populations (e.g., Kirkpatrick and Lande, 1989; Pál and Miklós, 1999; Bonduriansky and Day, 2009; Mullon and Lehmann, 2017), or under random transmission combined with limited gene flow (i.e., assuming that transmission within groups or spatial clusters that include parents and their offspring occurs randomly, Brown and Hastings, 2003; Hui et al., 2004; Silver, M and Di Paolo, E, 2006; Wakano, 2007; Lehmann, 2008; Han et al., 2009; Best et al., 2010; Débarre et al., 2012; Horns and Hood, 2012; Lion and Gandon, 2015; Mullon and Lehmann, 2018; Joshi et al., 2020; but see Ohtsuki et al., 2017, for a specific model of biased cultural inheritance under limited dispersal).

To fill this gap, we compute the selection gradient acting on a genetic locus with extended effects (e.g., on nest quality, pathogen load or cultural information) in a patch-structured population, where dispersal among patches is limited and extended effects can be transmitted across generations in a biased manner within patches. By disentangling and quantifying the various ways that a gene and its extended effects can be associated in such a scenario, our framework helps understand the nature of adaptation via non-genetic modifications. To illustrate this, we apply our framework to a model of gene-culture coevolution in which a genetically determined learning strategy coevolves with knowledge about the environment (e.g. Feldman and Cavalli-Sforza, 1976; Cavalli-Sforza and Feldman, 1981; Lumsden and Wilson, 1981; Boyd and Richerson, 1985; Aoki, 1986; Feldman and Laland, 1996; van Schaik, 2016). We show that the evolution of learning and the concomitant amount of knowledge generated by this evolution depends critically on the interaction between the degree of bias for cultural transmission within groups and the level of dispersal among groups. Finally, we discuss how our framework can be useful to study other biological problems, such as host evolution to pathogens, symbiotic mutualism, and niche construction.

## 2 Model

### 2.1 Population and traits

We consider a population of haploids subdivided among a large (ideally infinite) number of patches, all of size *n* (i.e., Wright’s island model). The population follows a discrete-time life-cycle consisting of three stages: adult reproduction;offspring dispersal;and competition among offspring to replace adults (see Figure 1a). Generations are non-overlapping but we allow for interactions between adults and offspring within a generation. Each individual in the population is characterised by a quantitative genetic character (e.g., breeding value for nest building, resistance to pathogen, social learning strategy) denoted by 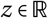 (see Table 1 for a list of symbols) and an extended trait 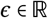 that can be transmitted between generations through ecological inheritance (e.g., nest quality, pathogen load, adaptive information about the environment; see section 4 for a specific example).

**Figure 1:**
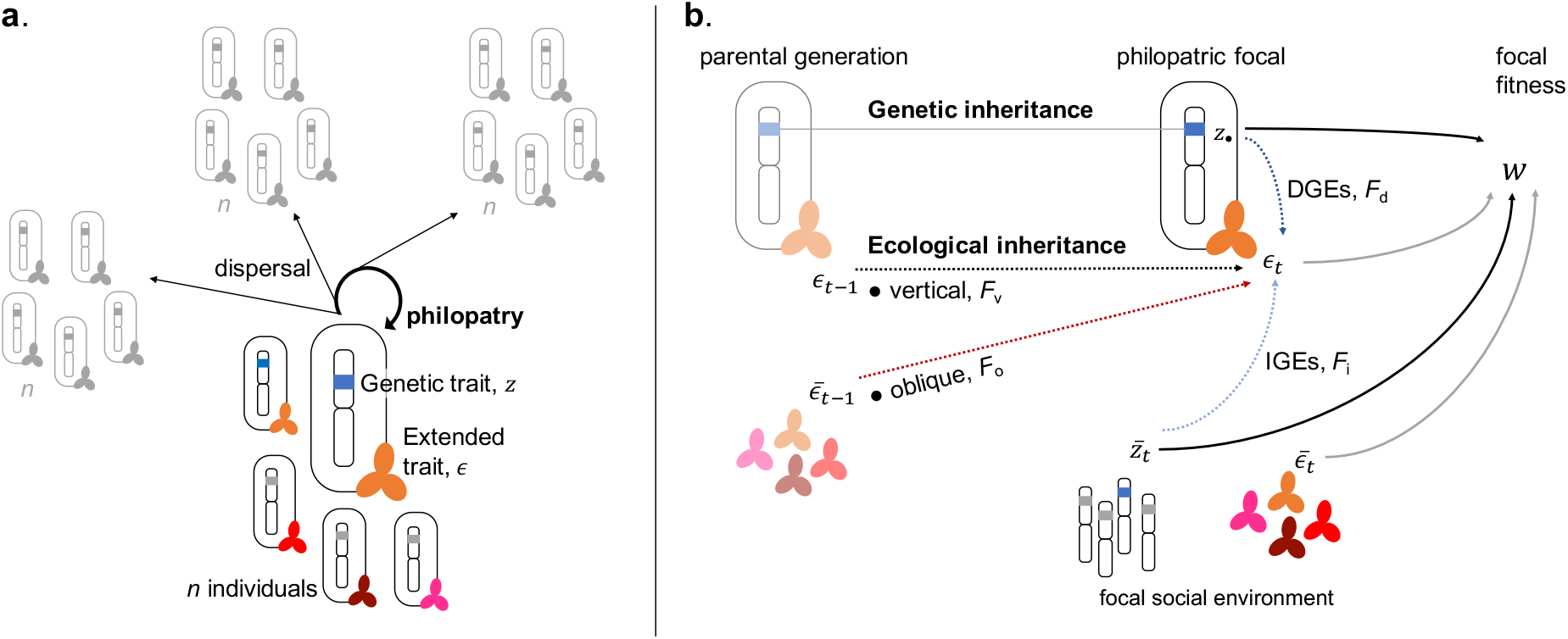
Genetic and extended traits under limited gene flow: pathways of inheritance, interactions and fitness effects. **(a)** The population consists of patches each home to *n* individuals. Every individual in the population is characterized by the genetic and extended traits it carries. Both traits can influence any step of the life cycle, which is as follows: (1) First, adults reproduce (in sufficient numbers to ignore demographic stochasticity);(2) Independently of one another, offspring either remain in their natal patch or disperse to a randomly chosen patch (so there is no isolation-by-distance);(3) Adults die and offspring compete locally for *n* open breeding spots in each patch. **(b)** While genetic traits are always inherited from parent to offspring, extended traits can be transmitted both: from the genetic parent when offspring remain in their natal patch (vertically) and from other individuals in the parental generation (obliquely). Once inherited, the extended trait of an individual can be modified directly by its own genetic trait (DGEs) and by the genetic trait of its neighbours (IGEs). Finally, the individual fitness of an individual depends on its own genetic and extended traits, as well as those expressed by neighbors in its patch.

**Table 1:** Main symbols and their meaning used in the general framework. This lists the main symbols and their meaning as used in sections 2 and 3.^†^ Relevant equations if applicable.

| Symbol | Meaning | Eq.<sup>†</sup> |
| --- | --- | --- |
| $n$ | Patch size. | |
| $m$ | Dispersal probability. | |
| $z$ | Quantitative genetic trait ($z_{\bullet}$, trait of a focal individual; $\bar{z}_t$, patch average at generation $t$; $z^m$, trait of a rare mutant). | |
| $\epsilon$ | Quantitative extended trait ($\epsilon_t$, of a focal individual at generation $t$; $\bar{\epsilon}_t$, patch average at generation $t$). | |
| $\hat{\epsilon}$ | Equilibrium extended trait in a population monomorphic for genetic trait $z$. | (3) |
| $F$ | Mapping for the modifications of the extended trait from one generation to the next. | (1)-(2) |
| $F_v(z)$ | Effect of vertical ecological inheritance on the extended trait of a philopatric individual in a population monomorphic for $z$. | (4) |
| $F_o(z)$ | Effect of oblique ecological inheritance on an individual's extended trait in a population monomorphic for $z$. | (4) |
| $F_E(z)$ | Total effect of ecological inheritance ($F_E(z) = F_v(z) + F_o(z)$). | |
| $F_d(z)$ | Direct extended genetic effects in a population otherwise monomorphic for $z$. | (5) |
| $F_i(z)$ | Indirect extended genetic effects in a population otherwise monomorphic for $z$. | (5) |
| $F_G(z)$ | Total extended genetic effects ($F_G(z) = F_d(z) + F_i(z)$). | |
| $w$ | Fitness of a focal individual. | (6) |
| $\bar{r}$ | Average within-patch relatedness (i.e., probability that two genes sampled at random with replacement within a patch are identical-by-descent). | (9) |
| $s(z)$ | Selection gradient on the genetic trait. | (7),(8) |
| $s_g(z)$ | Selection on non-extended genetic effects. | (8),(9) |
| $s_f(z)$ | Selection on extended genetic effects. | (8),(10) |
| $\mathcal{E}(z)$ | Extended genetic effect on carriers of a genetic mutation (decomposed as within- and trans-generational effects, $\mathcal{E}_W(z)$ and $\mathcal{E}_B(z)$, respectively). | (10),(11) |
| $\bar{\mathcal{E}}(z)$ | Extended genetic effect on the average non-genetic trait in the patch of carriers of a genetic mutation (decomposed as within- and trans-generational effects, $\bar{\mathcal{E}}_W(z)$ and $\bar{\mathcal{E}}_B(z)$, respectively). | (10),(16) |

### 2.2 Extended genetic effects and ecological inheritance

#### 2.2.1 Extended genetic effects

Within a generation, the extended trait of a focal individual can be influenced by genes via two pathways. First, it can be modified by the genetic character expressed by this focal (direct genetic effects, DGEs, Figure 1b); and second, by the characters of other patch members of the same generation (indirect genetic effects, IGEs, Figure 1b). These direct and indirect extended genetic effects capture how interactions between individuals of the same generation affect extended traits. For instance, if the character *z* is the production of extra-cellular antimicrobial agents, then the pathogen load of an individual, which in our context is its extended trait *ϵ*, depends on its own production and that of its patch neighbours.

#### 2.2.2 Ecological inheritance

Extended traits can be transmitted between individuals across generations via ecological inheritance (Figure 1b). We assume that such ecological inheritance occurs after dispersal. This assumption is natural when the extended trait consists of material resources that are physically tied to the patch so that an individual cannot disperse with it. For extended traits that can disperse with their carriers and are transmitted via social interactions, such as pathogens or cultural knowledge, our assumption entails that these interactions take place primarily after dispersal (we discuss this assumption at greater length in the Discussion). To specify ecological inheritance further in terms of vertical and oblique transmission, we consider philopatric and immigrant individuals separately below.

#### 2.2.3 Philopatric individuals

The extended trait of a philopatric individual can be transmitted from individuals of the previous generation both: (1) vertically from its genetic parent (vertical ecological inheritance, Figure 1b); and (2) obliquely from other individuals of the parental generation present in the patch (oblique ecological inheritance, Figure 1b). This distinction between vertical and oblique ecological inheritance allows to capture biased transmission due to non-random interactions within patches. The extended trait *ϵ_t_* of a focal philopatric individual at a generation *t* can thus be written as a function *F* of four variables,

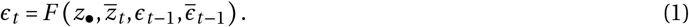

These four variables are: the genetic character of the focal individual, *z*_•_; the average genetic character in its patch at generation *t*, 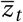; the extended trait of its genetic parent, *ϵ*_*t*−1_ (that lived at generation *t* − 1);and the average extended trait in the parental generation within its patch, 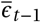 (see eqs. 20–22 for an explicit example of such a function *F*)*.

#### 2.2.4 Immigrants

If an offspring disperses, its genetic parent is absent from the patch it immigrates into. We assume that in the absence of family connections, an offspring interacts at random with adults of the previous generation. The extended trait of an immigrant individual at generation *t* is then given by

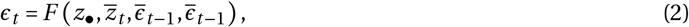

where *F*’s third argument is now the average extended trait in the parental generation in the patch the focal individual has immigrated into (instead of parental extended trait in eq. (1) for philopatric individuals).

#### 2.2.5 Trans-generational transformations of extended traits

The combination of modifications within generations and ecological inheritance can lead to cumulative carryover effects across generations, whereby individuals inherit modified extended traits that are then further modified and in turn transmitted to the next generation. Such dynamics, which are given by eqs. (1)–(2), unfold even in the absence of genetic evolution (see fig. 2b-c for e.g.). For our analysis, we assume that in the absence of genetic variation these dynamics do not lead to the unlimited transformation of the extended trait. In fact, we assume that in a genetically monomorphic population (so when all individuals have the same genetic trait *z*), the dynamics of the extended trait converge to an equilibrium 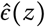. This equilibrium, which depends on the genetic character *z* but which we write as 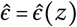 for short, must then satisfy

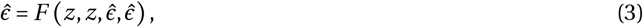

as well as the stability condition of 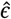:

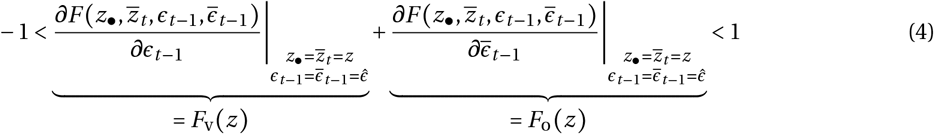

where *F*_v_(*z*) captures the effect of vertical ecological inheritance on the extended trait of a philopatric individual over one generation;and *F*_o_(*z*), the effect of oblique ecological inheritance. In particular, when ecological inheritance is random within a patch, then *F*_v_(*z*) = 0. Otherwise, effects of biased transmission between parents and offspring occur when *F*_v_(*z*) ≠ 0. As it will prove useful later, we also introduce the following notation to capture extended genetic effects within a generation,

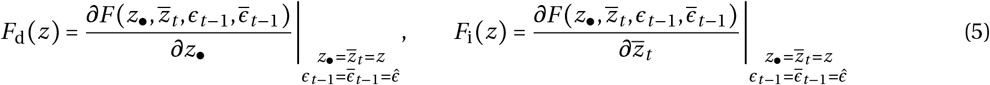

where *F*_d_(*z*) and *F*_i_(*z*) respectively measure the direct and indirect extended genetic effects (i.e., *F*_d_(*z*) is the marginal effect of an individual changing its genetic trait on its own extended trait, and *F*_i_(*z*), the effect of a change in the average genetic trait in the patch).

**Figure 2:**
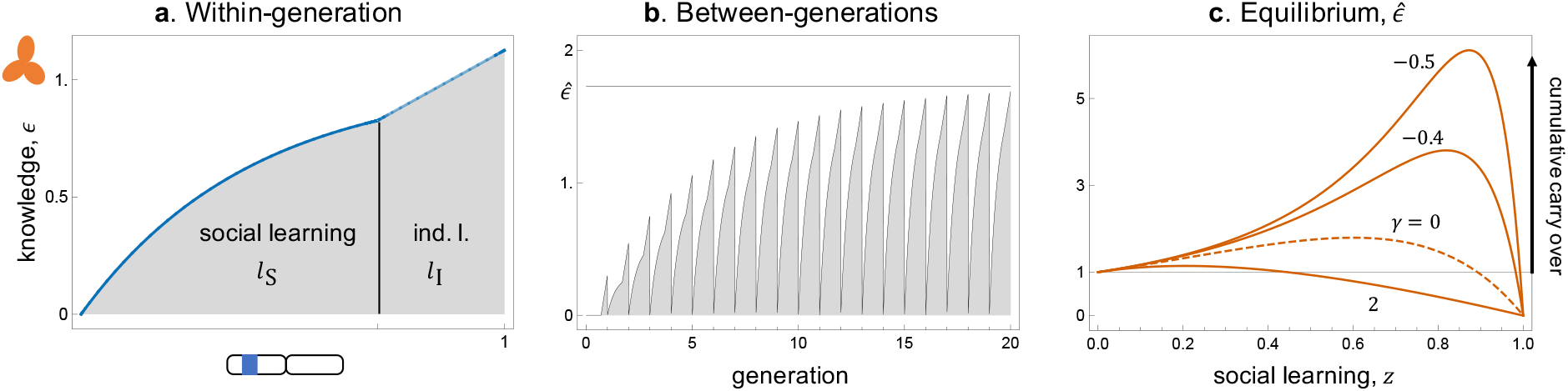
Within- and between-generations transformations of cultural information and its equilibrium. **(a)** Knowledge accumulation by one individual during its own life time, learning socially first according to *l*_S_ (eq. (20), with 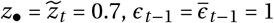, *β* = 2.5, *γ* = 0), and then individually according to *l*_I_ eq. (21), with *z*_•_ = 0.7, *α* = 1). **(b)** Knowledge accumulation across generations in a monomorphic population converging to equilibrium 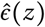 (found by iterating map *F*, eq. (22), starting with 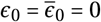, with 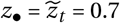, *β* = 2.5, *α* = 1, *γ* = 0). **(c)** Equilibrium knowledge in a monomorphic population according to investment into social learning *z* (given by eq. (25), with *β* = 2.5, *α* = 1, *γ* = −0.5, −0.4,0,2). As *α* = 1 here, the maximum knowledge that an individual can obtain in its lifetime via individual learning alone is 1. Any strategy that generates knowledge above 1 therefore entails cumulative culture. The learning strategy that generates maximum knowledge when expressed in the whole population is denoted by *z*_MAX_ (see eq. (26)).

### 2.3 Individual fitness

We assume that individuals with different combination of genetic and extended traits have different reproductive success. Specifically, the fitness *w* of an individual (defined as its expected number of successful offspring produced over one iteration of the life-cycle) depends on its genetic and extended traits, as well as those carried by its patch neighbours (fig. 1b). To capture this, we write the fitness of a focal individual at generation *t* with genetic and extended traits, *z*_•_ and *ϵ_t_* respectively, as a function

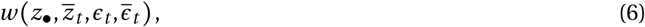

where 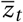 and 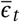 are the average genetic and extended traits, respectively, in the patch of the focal individual.

### 2.4 Evolutionary dynamics

To investigate the genetic and concommitant non-genetic evolution of our population, we derive the selection gradient, *s*(*z*), on the genetic trait *z*. This gradient gives the direction of selection, and thus information on the gradual evolution of *z* and its effect on extended trait *ϵ*. Specifically, the selection gradient determines singular genetic strategies (i.e., trait values *z*\* such that *s*(*z*\*) = 0) and their convergence stability (i.e., whether these singular strategies will be approached due to selection and the rare input of mutations with weak effects – when *s*′(*z*\*) < 0 – or not – when *s*′(*z*\*) > 0), Rousset, 2004; Dercole and Rinaldi, 2008). Equilibrium eq. (3) in turn allows investigating the extended trait expressed at such strategies and thus extended transformations concomitant to genetic evolution.

The selection gradient can be computed as the marginal change in the basic reproductive number, *R*_0_(*z*^m^, *z*), of a rare mutant with genetic trait *z*^m^ in a resident population that is otherwise monomorphic for genetic trait *z*,

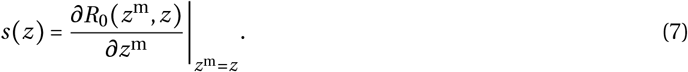

In the island model, this reproductive number is defined as the expected number of successful offspring produced by an individual that is randomly sampled from a local lineage (i.e., a lineage of individuals that reside in the same patch) of rare mutants with genetic trait *z*^m^ in a resident population with genetic trait *z* (Mullon et al., 2016; Lehmann et al., 2016, see Appendix A for details). Although the selection gradient can be straightforwardly computed numerically for a given model using eq. (7), our goal here is to unpack selection in a biologically meaningful way.

## 3 Selection gradient

We show in Appendix B that the selection gradient on a genetic trait with extended effects can be partitioned as the sum of two terms,

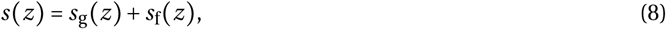

where the first, *s*_g_(*z*), is due to genetic effects on fitness only (i.e., ignoring extended genetic effects), while the second, *s*_f_(*z*), is due to extended genetic effects and how such effects feedback on fitness (see eqs. (B-8)–(B-9) for general expressions).

### 3.1 Selection due to (non-extended) genetic effects on fitness

The first component of selection is in fact given by the standard selection gradient on traits with fitness effects only (Frank, 1998; Rousset, 2004, for textbook treatments), i.e.,

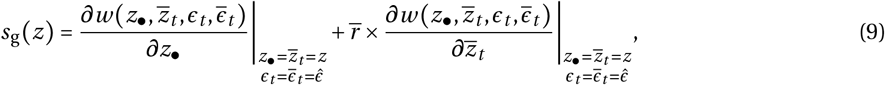

where 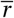 is average within-patch relatedness (here the probability that two individuals randomly sampled with replacement within the same patch carry an allele that is identical-by-descent at a neutral locus, see eq. (B-12)). Eq. (9) gives the standard decomposition of selection in subdivided populations, as a weighted sum of two genetic effects on fitness: the first is the direct effect of a focal individual changing its genetic trait on its own fitness; whereas the second is the indirect effect of a change in the focal on the fitness of an average patch member, weighted by average relatedness.

### 3.2 Selection due extended genetic effects and their feedback on fitness

Selection on extended genetic effects, meanwhile, is given by

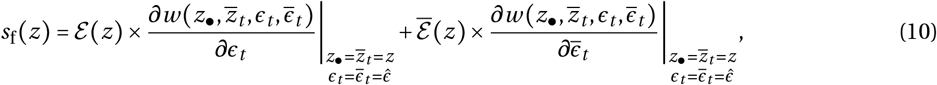

where 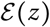 is the effect of a genetic change on the extended trait expressed by a representative carrier of this change, and 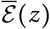 the effect of a genetic change on the average extended trait expressed by members of the patch in which a representative carrier of this change resides (in this context, “representative” refers to an average carrier of a rare genetic variant or mutation, where the average is taken over all possible genetic fluctuations that can occur within a patch;see eqs. (B-14)–(B-15) for mathematical definition). More intuitively, eq. (10) reflects the broad notion that evolutionary feedbacks via extended traits can occur in two non-exclusive ways: (1) a carrier of a mutation may express a different extended trait compared to non-carriers (difference whose magnitude is 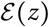), and this difference feeds back on the fitness of carriers (according to the first fitness derivative in eq. (10), which corresponds to the effect of a change in the extended trait of an individual on its own fitness); (2) carriers may reside in patches in which individuals on average express a different extended trait compared to individuals in other patches (with magnitude 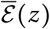), and it is this difference in social environments that in turn feeds back on the fitness of carriers (according to the second fitness derivative in eq. (10), which measures the effect of a change in the patch-average extended trait on the fitness of a member of that patch). We specify below how these two evolutionary feedbacks depend on modifications to the extended trait via genetic effects within generations and ecological inheritance between generations.

#### 3.2.1 Intra- and inter-generational extended genetic effects on a carrier

As an individual can influence its own extended trait, that of its current patch neighbours, as well as that of downstream individuals through ecological inheritance, we find that we can decompose the effect 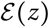 of a genetic change on the extended trait expressed by a representative carrier of this change depends on within-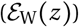 andbetween-generations effects 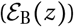,

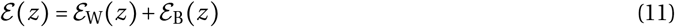

(see Appendix C for derivation). The intra-generational term simply consists of

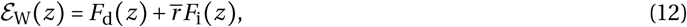

i.e., of the effect, *F*_d_(*z*), that a carrier of a genetic change has on its own extended trait, and on current relatives living in its patch, 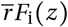. But due to ecological inheritance and limited gene flow, an individual may also influence the extended trait of downstream philopatric relatives. Selection owing to this process turns out to be

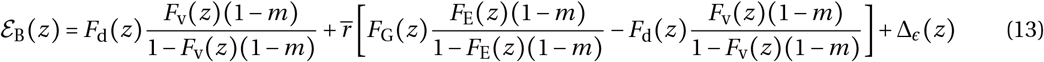

where *m* is the backward probability of dispersal (i.e., the probability that an individual is an immigrant in the absence of selection);*F*_G_(*z*) = *F*_d_(*z*) + *F*_i_(*z*), is the total extended genetic effects;*F*_E_(*z*) = *F*_v_(*z*) + *F*_o_(*z*), is the total effect of ecological inheritance; and

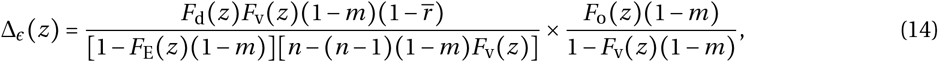

is due to the effect of random local genetic fluctuations on the extended trait (see eq. (C-42) in Appendix C for how we obtain this decomposition). Because in our later application (section 4) this latter term Δ_*ϵ*_(*z*) only influences trait evolution quantitatively and not qualitatively (not shown), we focus our attention on the rest of eq. (13), which also connects more easily to existing results.

The first term in eq. (13) is the effect that an individual has on its own extended trait, *F*_d_(*z*), and how this modification affects the extended trait of downstream philopatric descendants via vertical transmission. To see this, we can unpack said term as

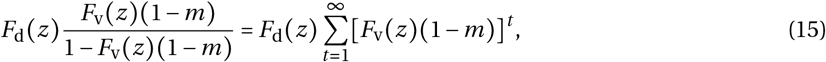

where the sum accumulates the direct extended genetic effects *F*_d_(*z*) originating from one individual (at time *t* = 0) across downstream philopatric generations (*t* = 1 onward) via vertical transmission. In the limiting case *m* = 0, eq. (15) in fact reduces to the trans-generational effects under selection calculated for a well mixed population under vertical transmission (see eqs. 15–17 of Mullon and Lehmann, 2017).

With limited gene flow (0 < *m* < 1), an individual can also influence the extended traits of downstream relatives by first influencing the extended traits of its patch neighbours, which are then transmitted across generations via ecological inheritance. Selection on such trans-generational effects is captured by the second term of eq. (13) (where the term within square brackets consists of the difference between the effects of all trans-generational modifications originating from one individual and those that are specifically due to its direct extended genetic effects transmitted vertically).

#### 3.2.2 Intra- and inter-generational extended genetic effects on the patch of a carrier

When the fitness of an individual also depends the extended trait of its neighbours, feedback selection can also occur via effect on the average extended trait in the patch of carriers (second term of eq. (10)). These effects can also be decomposed according to whether they occur within- or between-generations,

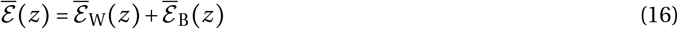

(see Appendix C for derivation), with intra-generational effects simply,

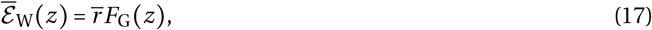

i.e., the total extended genetic effects of an individual weighted by average relatedness. The trans-generational transformation of the average extended trait, meanwhile, can be expressed as,

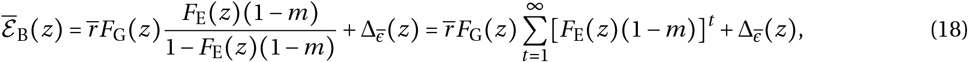

where

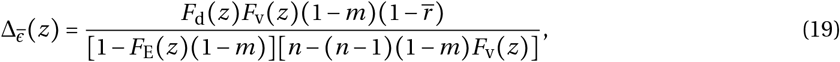

is again a term due to stochastic local genetic fluctuations. The first term of eq. (18) consists of the product between extended genetic effects among current relatives, 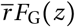, and how such effects impact the extended trait expressed by the downstream philopatric descendants of all these relatives via ecological inheritance (see right hand side eq. (18) for expansion of trans-generational effects into a sum). Eq. (18) aligns with previous analyses of selection on trans-generational effects that influence the condition of all individuals in a group equally (specifically, in the absence of preferential direct genetic effects – *F*_d_(*z*) = 0 – and vertical transmission – *F*_v_(*z*) = 0 – each individual in a group expresses the same extended trait, then eq. (18) reduces exactly to eq. (29) of Mullon and Lehmann, 2018; see also eq. (9) of Lehmann, 2007 and eq. (4.11) of Sozou, 2009 for similar expressions).

By contrast to these previous studies, our model allows for differential expression of the extended trait within a patch due to direct genetic effects and/or vertical ecological inheritance. Our extension is thus especially relevant to understand evolutionary driven changes in individual condition, such as micro-environments (e.g., nests or burrows), pathogens or microbiome, cellular state or culture. As we have shown, selection in this case depends on multiple feedbacks on the fitness of relatives (eqs. (8)–(10)). This is because a carrier of a genetic mutation not only modifies (i) its own extended trait, but also (ii) the extended trait of individuals it interacts with during its lifetime via indirect effects, as well as (iii) the extended trait of individuals in downstream generations via ecological inheritance (see fig. 1b). Due to limited gene flow, these other affected individuals are either carriers of the genetic mutation (i.e., relatives) or non-carriers that interact with relatives. In either case, the modifications initiated by a carrier of a genetic mutation feeds back on the fitness of current and downstream relatives. Our analysis disentangles the various pathways via which such evolutionary feedbacks occur, owing to direct (*F*_d_) and indirect (*F*_i_) extended genetic effects combined with vertical (*F*_v_) and oblique (*F*_o_) ecological inheritance (eq. (11)–(18)). As we show in the next section by applying our framework, this decomposition can help understand how natural selection shapes genetic traits and the modifications these entail.

## 4 Gene-culture coevolution under limited gene flow

To illustrate our general result, we investigate a model of gene-culture coevolution, whereby a genetically determined learning behavior co-evolves along culturally transmitted information (Feldman and Cavalli-Sforza, 1976; Lumsden and Wilson, 1981; Boyd and Richerson, 1985; Aoki, 1986; Feldman and Laland, 1996; van Schaik, 2016).

### 4.1 Assumptions

We assume that after dispersal, offspring acquire adaptive culture or information (e.g., foraging skills). They acquire such information via two routes: first, they learn socially from the adults in their patch (e.g., by imitation) and second individually (e.g., by trial and error). The evolving genetic trait is the investment 0 ≤ *z* ≤ 1 of time or energy into social learning (so that 1 – *z* is invested in individual learning) and the extended trait *ϵ* ≥ 0 is the amount of knowledge held by an individual. The combination of social and individual learning allows for the accumulation of knowledge across generations, i.e., cumulative culture, which is thought to be a hallmark of human populations (Boyd et al., 2011; van Schaik, 2016). If a body of theoretical literature has helped better understand the conditions that favour cumulative culture, most models assume that populations are well mixed and/or that individuals learn socially from one another at random (Boyd and Richerson, 1995; Enquist et al., 2007; Borenstein et al., 2008; Aoki et al., 2012; Lehmann et al., 2013; Nakahashi, 2013; Wakano and Miura, 2014; Aoki and Feldman, 2014; Kobayashi et al., 2015; Mullon and Lehmann, 2017; but see Rendell et al., 2010; Ohtsuki et al., 2017; Kobayashi et al., 2019). Here, we use our framework as a platform to investigate the evolution of learning strategies and cumulative culture under the joint effects of limited gene flow among groups and non-random social learning within groups. As a further extension to previous models, we also allow for social learning by different individuals to interact with one another in a synergistic or antagonistic manner. Synergy could for instance occur when social learners help one another while antagonism could arise where gathering adaptive information from the social environment is competitive.

#### 4.1.1 Cultural dynamics

##### Social learning

In terms of within generation cultural dynamics, an offspring is born with zero knowledge and after dispersal, first learns socially from the adults present in the patch. A focal offspring accumulates knowledge via this route in a way that decelerates with the amount of investment 0 ≤ *z*_•_ ≤ 1 made into social learning and plateaus to the level of knowledge carried by present adults (fig. 2a), reflecting that an offspring cannot gain more knowledge than the individuals it learns from. We allow for philopatric offspring to preferentially learn vertically from their parent, with weight 0 ≤ *υ* ≤ 1, compared to a random adult (with weight 1 – *υ*, see Table 2 for list of symbols used specifically for the gene-culture coevolution model). The parameter *υ* controls how biased transmission is towards parents and thus how vertical cultural transmission is compared to oblique (so influencing *F*_v_ and *F*_o_, fig. 1b). We also allow for the efficiency with which an individual learns socially to depend on the investment of other patch members into social learning according to a parameter *γ*. Specifically, we assume that the amount of knowledge obtained socially by a philopatric individual investing *z*_•_ into social learning at generation *t* when its patch neighbours have invested on average 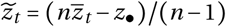, is

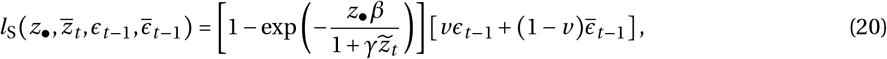

where the first term within square brackets is the effective transmission efficiency of social information and the second term in square brackets is the maximum target knowledge that can be socially transmitted (which is a weighted average of parental knowledge in the patch). In eq. (20), the parameter *γ* > −1 controls interference among social learners: when *γ* = 0, there is no interference and social information is transmitted with baseline efficiency *β* (as in e.g., Lehmann et al., 2013; Wakano and Miura, 2014; Kobayashi et al., 2015; Ohtsuki et al., 2017; Mullon and Lehmann, 2017);when *γ* < 0, social transmission is enhanced by other individuals;and when *γ* > 0, it is impaired. In the context of our general framework, the parameter *γ* therefore modulates the direction and strength of indirect extended genetic effects (so influencing *F*_d_ and *F*_i_, fig. 1b).

**Table 2:**
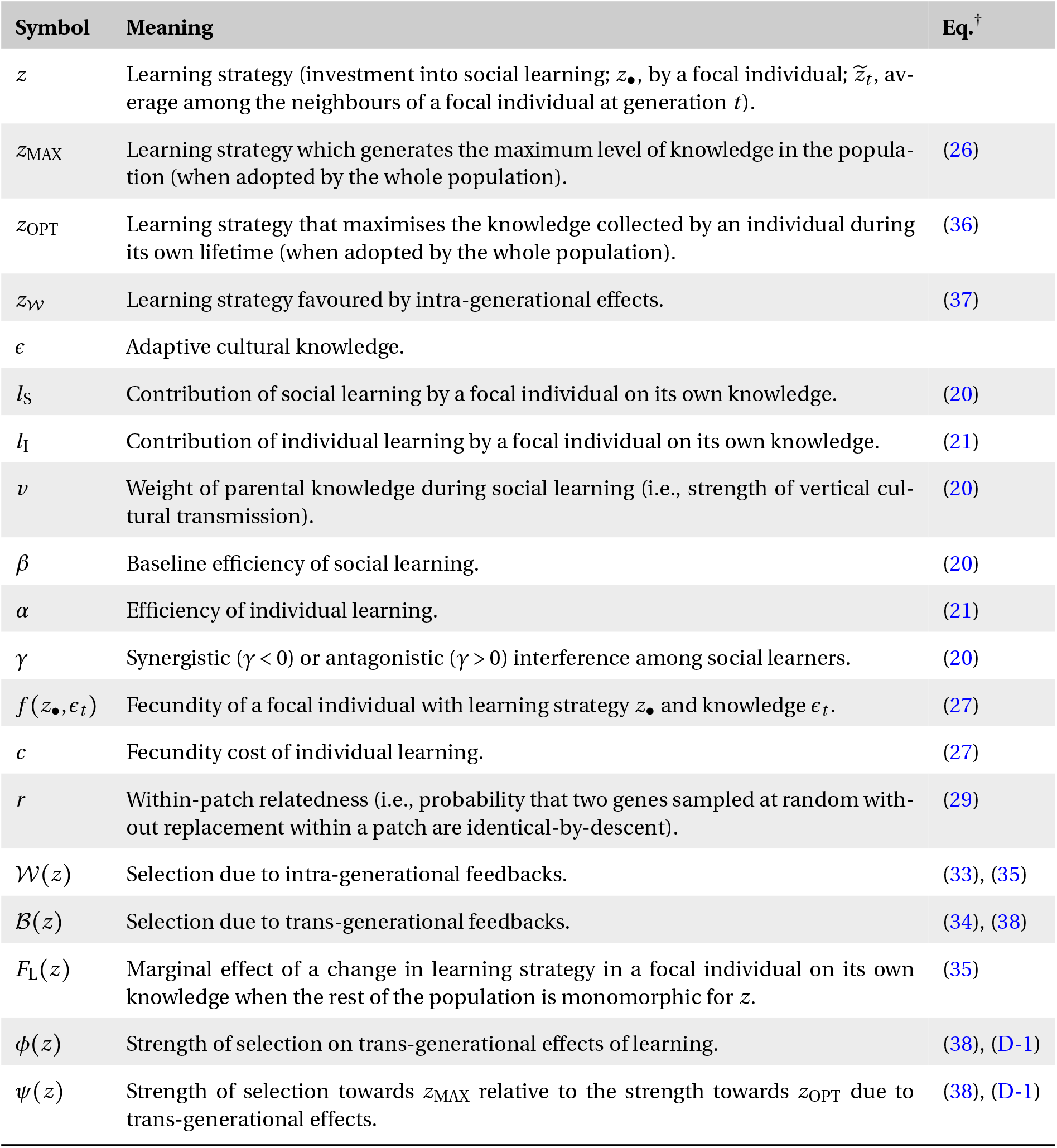
Main symbols and their meaning used in the gene-culture coevolution model. This lists the main symbols and their meaning as used in section 4.^†^ Relevant equations if applicable.

##### Individual learning

After learning socially, an offspring learns individually, and accumulates knowledge linearly with the investment, 1 – *z*_•_, into individual learning according to

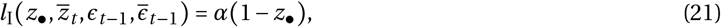

where *α* > 0 tunes the efficiency of individual learning (fig. 2a, as in e.g. Lehmann et al., 2013; Wakano and Miura, 2014; Kobayashi et al., 2015; Ohtsuki et al., 2017; Mullon and Lehmann, 2017). The knowledge that a philopatric individual has at the time of reproduction at generation *t* is then given by,

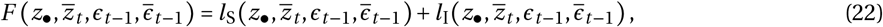

the sum of socially and individually acquired information,

##### Learning strategy under individual control that maximises knowledge

All else being held constant, the learning strategy that maximises the knowledge that a focal individual obtains is the strategy 0 ≤ *z*_•_ ≤ 1 such that,

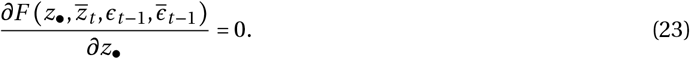

Substituting eqs. (20)–(22) into the above equation, we find that if it exists, this individual strategy can be written as,

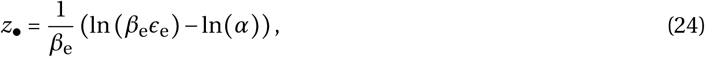

which increases with 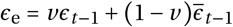, the amount of knowledge accessible to the focal individual (and also depends on 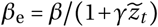, the effective rate of transmission via social learning). This amount *ϵ*_e_ depends on the knowledge carried by the parent and its neighbours, which in turn depends on their ancestors’ level of knowledge and so on.

##### Equilibrium cultural dynamics in a monomorphic population

In a population genetically monomorphic for learning strategy *z*, these cultural dynamics converge to an equilibrium,

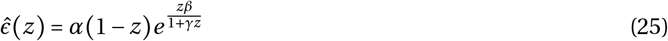

(found by substituting eqs. (20)–(22) into eq. 3);it is straightforward to show that this equilibrium is stable, i.e., that eq. 4 holds). Eq. (25) displays characteristic effects of social learning on culture (see also fig. 2b-c): knowledge initially increases with social learning (provided *β* > 1), leading to knowledge being accumulated across generations (i.e., individuals acquire more knowledge than they would have been able to by individual learning alone, 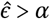), but past a threshold of social learning, knowledge decreases and eventually collapses as no individuals in the population produce knowledge via individual learning (i.e., 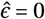 when *z* = 1). Interference among social learners (*γ* ≠ 0) does not change this relationship between equilibrium knowledge and learning strategy (fig. 2c), but knowledge reaches greater levels when social learning is synergistic (*γ* < 0) than when it is antagonistic (*γ* > 0).

The relationship between equilibrium knowledge and learning strategy in a monomorphic population (eq. (25), fig. 2c) implies that there exists a learning strategy such that if adopted by the whole population, generates the maximum possible level of knowledge. This optimal strategy, say *z*_MAX_, is determined by

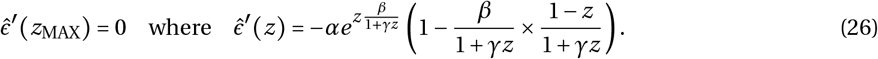

In the absence of interference (*γ* = 0) this strategy simply is *z*_MAX_ = 1 – 1/*β*, i.e., as social learning efficiency increases, more resources invested into social learning generate greater knowledge. Compared to this baseline, synergistic interactions among social learners (*γ* < 0) increase *z*_MAX_, and antagonistic interactions decrease it (eq. (26), fig. 2c). Whether selection favours the evolution of such an optimal strategy, however, depends on the fitness effects of learning and knowledge, which we describe next.

#### 4.1.2 Fitness effects

In terms of fitness, we assume that an individual’s fecundity increases with the amount of adaptive information it has collected but decreases with the amount of resources invested into individual learning. Social learning, by contrast, is assumed to be cost free for simplicity. These assumptions reflect the notion that social learning is cheap compared to individual learning as it outsources risk and helps avoiding fatal mistakes (Boyd and Richerson, 1985). One way to formalise this is to write the fecundity of an individual with information level *ϵ_t_* and social learning strategy *z*_•_ as a sum of these two factors,

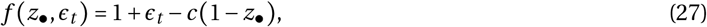

where *c* > 0 is a parameter tuning the cost of individual learning (we also explored multiplicative effects on fecundity and this did not influence our results qualitatively, not shown). The opposite effects of knowledge and individual learning on fecundity (eq. (27)), combined with the fact that knowledge ultimately breaks down when individual learning is absent in the population (eq. (25), fig. 2c), lead to a social dilemma: on one hand, individuals have an incentive to invest all their resources into social learning, but on the other, if every individual in the population does so, then there is no adaptive information to actually learn.

The fitness of a focal individual with fecundity *f*(*z*_•_, *ϵ_t_*) in the island model of dispersal is then given by,

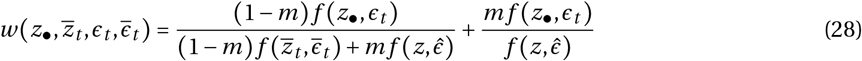

(when individuals produce a large – effectively infinite – number of zygotes, e.g., Rousset, 2004), where the first summand represents philopatric fitness (i.e., the expected number of offspring that secure a breeding spot in their natal patch), which is given by the ratio of the focal’s offspring that remain in their natal patch ((1 – *m*) *f* (*z*_•_, *ϵ_t_*) with *m* as the probability of dispersal) to the total number of offspring that enter competitionin this patch, consisting of all philopatric offspring 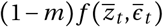^†^ and immigrants from other patches 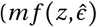 where *z* is the investment in social learning in other patches, which can be assumed to be monomorphic with resulting equilibrium knowledge 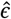 given by eq. (25));and the second summand of eq. (28) is dispersal or allopatric fitness (i.e., the expected number of offspring that secure a breeding spot in non-natal patches), which is the ratio of the focal’s offspring that emigrate to the expected total number of offspring in a non-natal patch.

To perform the analysis of selection elaborated in section 3, we further need to specify average within-patch relatedness, 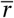 (see below eq. (9) for definition). This relatedness coefficient can be decomposed as

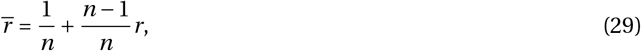

where *r* is the probability that two individuals randomly sampled without replacement within the same patch carry an allele that is identical-by-descent at a neutral locus. Such probability, which is connected to the classical notion of *F*_ST_ from population genetics (Rousset, 2002), can be derived from standard coalescence arguments, yielding

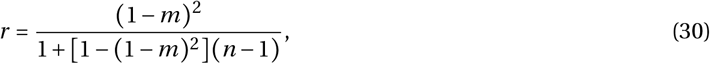

in the island model of dispersal with non-overlapping generations (e.g., Rousset, 2004).

### 4.2 Selection on social learning

#### 4.2.1 Genetic effects on fitness

To understand how selection shapes social learning and adaptive knowledge in our model, let us first investigate selection on social learning ignoring its extended effects on knowledge (so focusing on eq. (9)). Substituting eqs. (27)–(30) into eq. (9), we find that selection in this case is

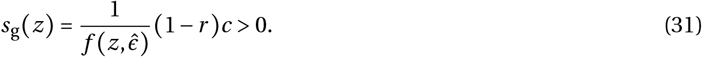

Because individual learning is more expensive than social learning (*c* > 0), this selection component is always positive (see red curve in fig. 3a), indicating that in the absence of feedback, selection always favours an increase in social learning, leading to individuals investing all their resources into social learning and none into individual learning (*z* = 1).

**Figure 3:**
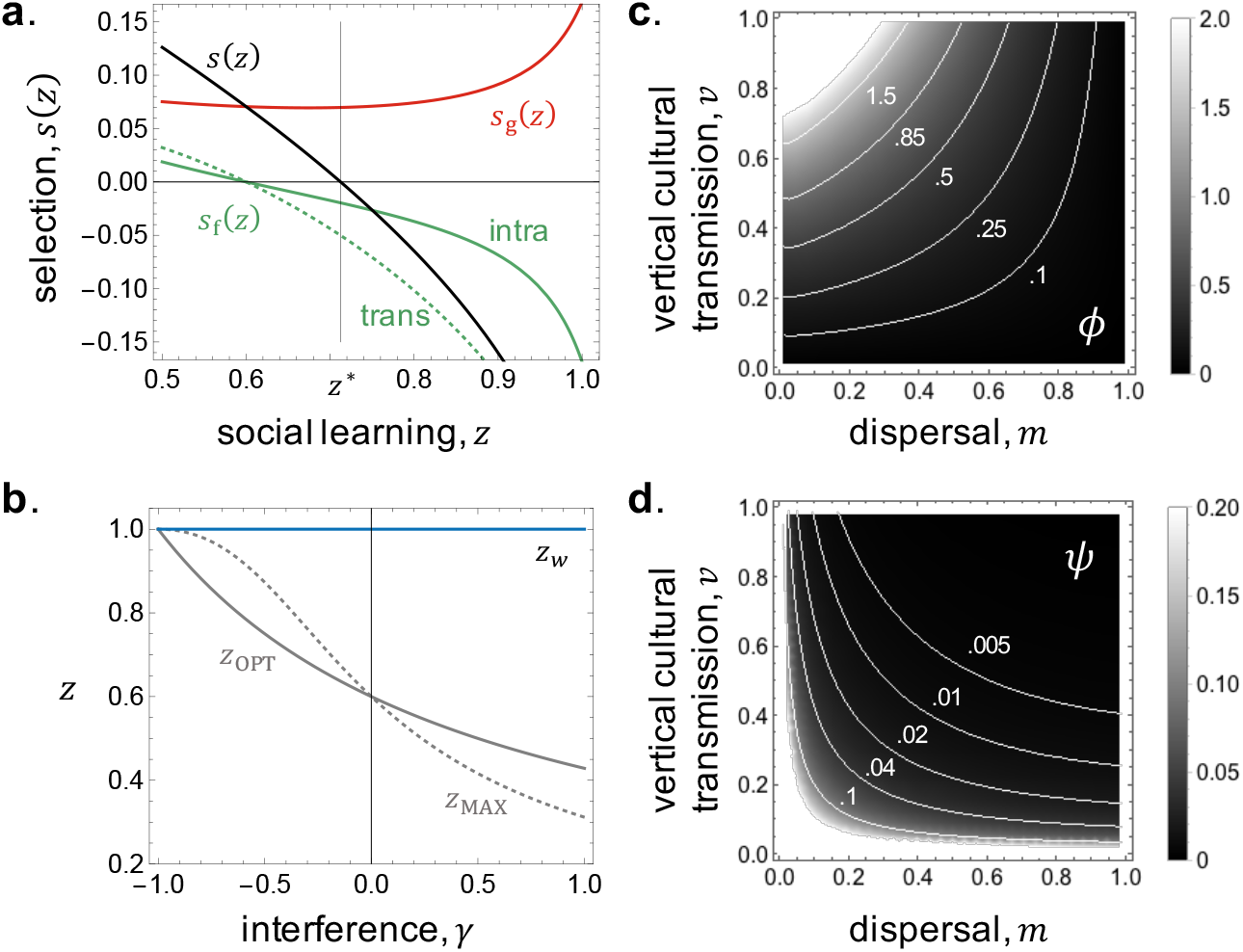
Selection on social learning. **(a)** Decomposition of the selection gradient *s*(*z*) (in black) according to genetic *s*_g_(*z*) (in red) and feedback effects *s*_f_(*z*) (in green), which are further decomposed into intra- (full line) and trans-generational (dashed line) effects (computed from eqs. (31)–(35) and eqs. (38) and (D-1), with *β* = 2.5, *c* = *α* = 1, *γ* = 0, *υ* = 1, *m* = 0.01, *n* = 10). The evolutionary convergent strategy *z*\* = 0.7 (where *s*(*z*\*) = *s*_g_(*z*\*) + *s*_f_(*z*\*) = 0) corresponds to a balance between selection due to genetic effects *s*_g_(*z*) (in red), which favours investing all resources into social learning, and feedback selection *s*_f_(*z*) (in green), which favours greater levels of individual learning. **(b)** Evolutionary convergent learning strategy in a well-mixed population as a function of interference parameter *γ* (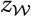 in blue, computed from eq. (37), same parameters as (a)). Also shown are the strategies that maximise adaptive information at the individual-(*z*_OPT_, full gray line, eq. (36)) and population-level (*z*_MAX_, dashed gray line, eq. (26)). We see that both of these strategies are equal in the absence of interference *γ* = 0. **(c)-(d)** The effect of vertical cultural transmission *υ* and dispersal *m* on: the strength of selection on trans-generational effects, *ϕ*(*z*) (in (c) from eq. (D-1) with *γ* = −0.5, *z* = 1;other parameters same as (a)), and on the strength of selection towards *z*_MAX_ relative to the strength towards *z*_OPT_ due to trans-generational effects, *ψ*(*z*) (in (d) from eq. (D-1) with *n* = 2;other parameters same as (c)). Lighter shade means greater values (see figure legend). This shows that selection *ϕ*(*z*) on trans-generational effects increases as dispersal becomes limited (*m* decreases) and cultural transmission becomes vertical (*υ* increases). Meanwhile selection term *ψ*(*z*) tends to promote the evolution of *z*_MAX_ rather than *z*_OPT_ when cultural inheritance is random (*υ* decreases) and relatedness is high within groups (*m* decreases, main text for interpretation).

#### 4.2.2 Feedback selection

Selection however also depends on the way that social learning influences knowledge and how this feeds back on the fitness of relatives (eq. (10)). Plugging eqs. (20)–(30) into eq. (10) (with eqs. (11)–(14), (16)–(19)), we find that selection due to such feedbacks can be partitioned as

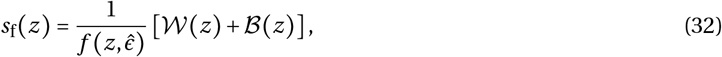

where

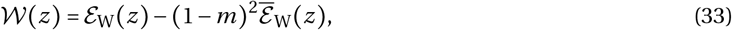

corresponds to selection due to intra-generational feedbacks, and

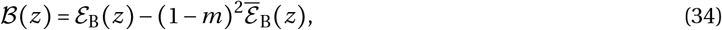

due to trans-generational feedbacks. At the broad scale described by eq. (33)–(34), selection on learning due to feedbacks depends on the effect that social learning by an individual has (i) on the knowledge of its current and downstream relatives in its patch (including itself, 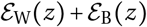), and (ii) on the average knowledge in its patch and experienced by its downstream relatives, 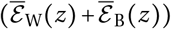, which is weighted by – (1 – *m*)^2^ owing to kin competition (because for e.g. when an individual increases adaptive information for all patch members, such an increase exacerbates competition for current and future relatives within the patch).

##### Intra-generational feedback

In terms of model parameters, selection due to intra-generational feedbacks reads as,

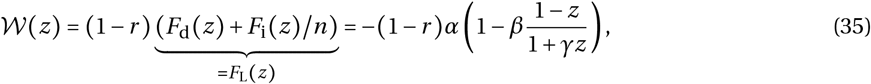

where *F*_L_(*z*) gives how a change in an individual’s learning strategy influences its own knowledge in a population otherwise monomorphic for *z*. As a result, the component of selection 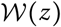 alone favours a combination of individual and social learning,

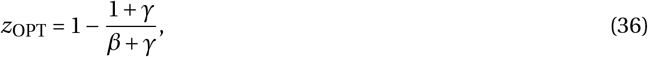

which when adopted by the whole population, maximises the level of adaptive information an individual collects within its own lifetime for itself (i.e., when the population is monomorphic for *z*_OPT_, any mutant will collect lower levels of adaptive information within its own lifetime – with *z*_OPT_ such that *F*_L_(*z*_OPT_) = 0, eq (35), see full green curve fig. 3a). In the absence of interference among social learners (*γ* = 0), this strategy also maximises knowledge in the entire population (i.e., *z*_OPT_ = *z*_MAX_, eq. (26), fig. 3b). When social learning is synergistic (*γ* < 0), however, the individual-strategy that maximises individual knowledge consists of less social learning than the population-strategy that maximises knowledge at the population-level (*z*_OPT_ < *z*_MAX_, fig. 3b). This is because the latter considers changes in learning strategy in all individuals, rather than just in a focal one. With synergy, all individuals performing more social learning generates more knowledge than when performed by a single individual, leading to *z*_OPT_ < *z*_MAX_. Conversely, when social learning is antagonistic (*γ* > 0), then *z*_OPT_ > *z*_MAX_ (fig. 3b).

##### Total intra-generational effects

If we add selection due to intra-generational feedback effects (eq. (32) with 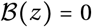) and selection due to genetic effects on fitness (which also occur within generations, eq. (31)), we obtain the selection gradient on social learning due its total intra-generational effects. Such selection favours the evolution of a learning strategy, 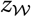, given by,

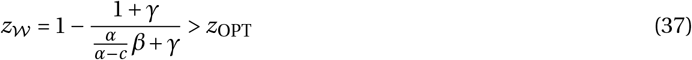

(i.e., such that 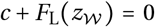, Fig. 3b). Under this strategy 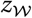, less resources are invested into individual learning than under *z*_OPT_ (eq. (36)) due to the fitness cost *c* of individual learning. Note that total selection on intra-generational effects favours the same learning strategy 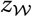 in a well-mixed and dispersal-limited population (i.e., eq. (37) does not depend on *m*). This is due to our assumptions that generations are non-overlapping and patches are of constant size, in which case the benefits from interacting with relatives are exactly offset by the cost of competing with them under limited gene flow (Taylor, 1992). Note also that selection on intra-generational effects is independent from the mode of cultural transmission (i.e., eq. (37) is independent from *υ*).

##### Trans-generational feedback

Owing to cultural inheritance and limited gene flow, however, feedbacks between relatives can also occur across generations. We find that selection due to such feedbacks can be expressed as

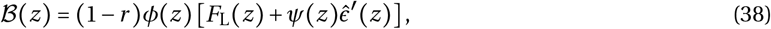

where *ϕ*(*z*) ≥ 0 and *ψ*(*z*) ≥ 0 are complicated non-negative functions of *z* and model parameters (see eq. (D-1) in appendix D for details). Inspecting the term within square brackets of eq. (38) reveals that selection due to trans-generational feedbacks is composed of two forces: one that favours the strategy *z*_OPT_ (according to *F*_L_(*z*));and another *z*_MAX_ (according to 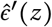). Both of these forces are proportional to *ϕ*(*z*), which can be interpreted as the strength of selection on trans-generational effects. The function *ψ*(*z*), meanwhile, characterises the strength of selection towards the strategy *z*_MAX_ relative to the strength towards *z*_OPT_ due to trans-generational effects.

To better understand the nature of selection on trans-generational effects, let us first consider a scenario where there is no interference among social learners, *γ* = 0. In this case *z*_OPT_ = *z*_MAX_ (Fig. 3b), so 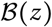 favours a single strategy that maximises knowledge both at the individual and population level, with strength proportional to *ϕ*(*z*) (eq. 38). Since 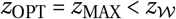 (Fig. 3b), selection due to trans-generational feedbacks favours less investment into social learning and more into individual learning compared to selection due to intra-generational effects (see green dashed curve Fig. 3a). In addition, numerical exploration of *ϕ*(*z*) reveals that selection on trans-generational effects increases as dispersal becomes limited and cultural transmission becomes vertical (i.e., *ϕ*(*z*) increases as *m* → 0 and *υ* → 1, Fig. 3c). We therefore expect that under these conditions, selection leads to greater investment into individual learning and greater levels of adaptive knowledge (in agreement with the results of Ohtsuki et al., 2017, who assumed that *γ* = 0). Intuitively, this is because as dispersal becomes limited and cultural transmission becomes vertical, the association between genetic and knowledge variation increases, so that the effects of a change in learning strategy are increasingly tied to individuals that express this change. As a result, the effects of learning on knowledge are increasingly apparent to selection.

When social learners interfere with one another (*γ* ≠ 0), however, the learning strategies that maximise knowledge at the individual and population level disagree (*z*_OPT_ ≠ *z*_MAX_). This raises the question: when does selection due to trans-generational effects favour the strategy *z*_MAX_ that leads to the greatest level of knowledge when expressed in the whole population? Numerical examination of *ψ*(*z*) shows that selection due to trans-generational effects tends to promote the evolution of *z*_MAX_ rather than *z*_OPT_ when cultural inheritance is random and relatedness is high within groups (i.e., *ψ*(*z*) increases as *m* and *υ* decrease, Fig. 3d). This can be understood by considering a rare mutant who invests more resources into social learning than a common resident who expresses *z*_OPT_, when social learning is synergistic (*γ* < 0, so that the mutant strategy is between *z*_OPT_ and *z*_MAX_). This change in strategy decreases the knowledge of the mutant (as strategy is different to *z*_OPT_) but increases the knowledge of contemporary neighbours due to synergy. In turn, when cultural transmission is purely vertical (*υ* = 1), this difference extends to descendants: philopatric offspring of the mutant receive less knowledge than other philopatric offspring. By contrast, when cultural transmission is random within patches (*υ* = 0), offspring of the mutant benefit from the increased knowledge of neighbours while other offspring suffer from learning poorer knowledge from the mutant. These mitigating effects of random transmission on the difference in knowledge between different offspring increase as dispersal becomes limited and as there are fewer adults in the patch. Accordingly, trans-generational effects then disfavour any strategy other than *z*_OPT_ when *υ* = 1 but favour strategies closer to *z*_MAX_ when *υ* = 0 and relatedness within patches is high.

At a superficial level, our analysis of *ψ*(*z*) suggests that in the presence of interference among social learners (*γ* ≠ 0), the evolution of learning leads to more knowledge when cultural inheritance is random rather than vertically biased. It is however important to keep in mind that *ψ*(*z*) is a relative measure of the strength of selection favouring *z*_MAX_ compared to *z*_OPT_ (eq. (38)). The overall strength of selection due to trans-generational effects is given by *ϕ*(*z*) (eq. (38)), which increases as cultural inheritance becomes vertically biased. Vertical cultural inheritance therefore has antagonistic effects on knowledge accumulation through learning evolution when *γ* ≠ 0: on one hand, it increases the relevance of trans-generational effects compared to intra-generational effects, but on the other it favours the evolution of strategies that do not maximise knowledge at the population level. We investigate the outcome of such antagonistic effects in greater depth in the next section.

#### 4.2.3 Evolutionary convergent strategies and cumulative culture

To investigate trans-generational effects further, we computed numerically the evolutionary convergent learning strategy, *z*\* (i.e., that towards which the population will converge under gradual evolution), which satisfies,

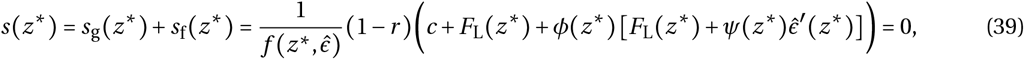

(found by adding eqs. (31) with (32) and using eqs. (35) and (38)), as well as the resulting level 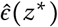 of knowledge such a strategy yields (using eq. (25)) for various model parameters (see Fig. 4). In the absence of interference among social learners (*γ* = 0), we find that individual learning is favoured when gene flow is limited (*m* is small) and cultural transmission is vertical (*υ* is large, Fig. 4a) which is in line with our analysis of eqs. (38) when *γ* = 0. In turn, the evolution of such learning strategies leads to the accumulation of greater levels of adaptive information in the population (Fig. 4b). Note that since selection due to intra-generational effects are independent from vertical cultural inheritance and dispersal (eq. (37)), the effects observed in Fig. 4 are entirely driven by trans-generational effects. Through its negative effects on trans-generational relatedness, a large group size tends to favour the evolution of social rather than individual learning leading to lower levels of adaptive information (Supplementary Figure 1). However, this negative effect of increased group size on the association between the learning strategy and knowledge of individuals is weak compared to the effect of increased dispersal or decreased vertical transmission. So provided some information is transmitted vertically and dispersal is limited, significant levels of information can accumulate in a population of large groups where genetic relatedness is low (Supplementary Figure 1).

**Figure 4:**
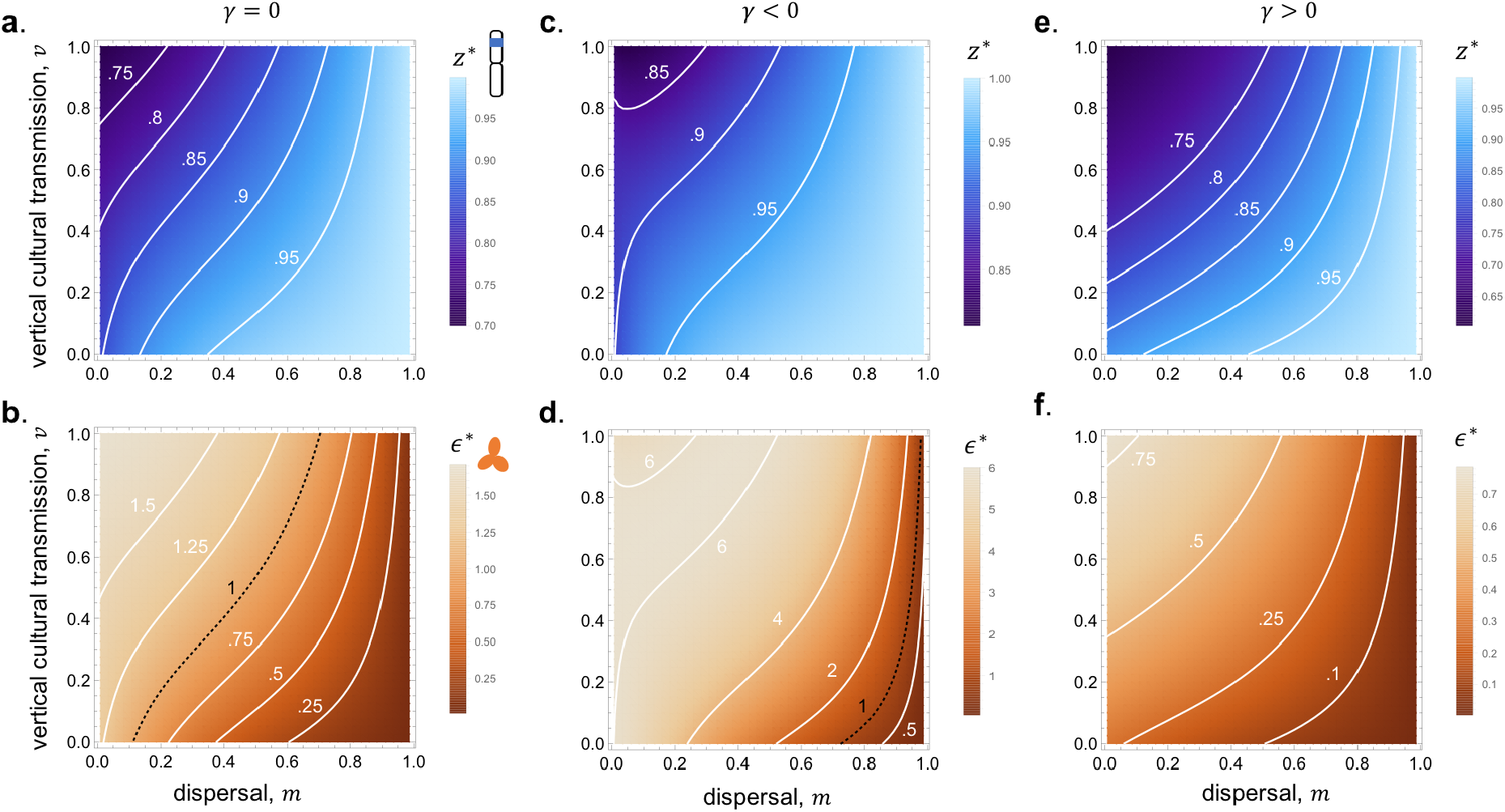
The influence of vertical cultural transmission, limited gene flow and social learning inter-ference on gene-culture coevolution. **Top row (a,c,e):** Contours of evolutionary convergent learning strategy *z*\* according to vertical cultural transmission on y-axis and limited gene flow on x-axis (com-puted by solving numerically eq. (39) for *z*\*, parameter values: *β* = 2.5; *c* = *α* = 1; *n* = 10; (a) *γ* = 0; (c) *γ* = −0.5; (e) *γ* = 2;lighter colours mean greater investment *z*\* in social learning – so less individual learning – see colour figure legend). These show that social learning (high *z*\*) is favored by high dispersal (small *m*), low vertical inheritance (high *υ*) and synergistic learning (*γ* < 0). **Bottom row (b,d,f)**: Contours of equilibrium knowledge 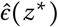 at evolutionary convergent learning strategy *z*\* according to vertical cultural transmission on y-axis and limited gene flow on x-axis (computed from values found in top row for *z*\* and eq. (25), same parameters as top row). Contour for 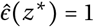, which is the threshold above which cumulative culture occurs (see Fig. 2c), shown as a black dashed line (lighter colours mean greater knowledge in the population, see colour figure legend). The conditions that favour the greatest level of adaptive knowledge are therefore low dispersal (small *m*), high vertical inheritance (high *υ*) and synergistic learning (*γ* < 0, main text for interpretation).

While synergistic learning (*γ* < 0) favours greater investment into social learning (Fig. 4c) that can nonetheless lead to high levels of cumulative knowledge (Fig. 4d), antagonistic learning (*γ* > 0) favours greater individual learning (Fig. 4e) but engenders low levels of non-cumulative knowledge (Fig. 4f). In agreement with our analysis of the antagonistic effects of cultural inheritance bias on knowledge accumulation through learning evolution, we observe that knowledge in the population is not maximised by full vertical cultural inheritance in the presence of synergistic social learning, *γ* < 0 (see top left corner of Fig. 4d). Rather, the accumulation of knowledge is greatest for intermediate levels of vertical cultural inheritance and gene flow, and these critical levels are independent from group size as long as it is moderate (*n* ≳ 10 for our parameters, Fig. 5). Such intermediate levels of inheritance and gene flow ensure that inter-generational effects are strong enough to offset the costs *c* of individual learning, but not so strong that they lead to investing so many resources into individual learning that knowledge in the population decreases (i.e., intermediate levels ensure that *c* + (1 + *ϕ*(*z*_MAX_))*F*_L_(*z*_MAX_) = 0, where *z*_MAX_ is given by eq. (26)).

**Figure 5:**
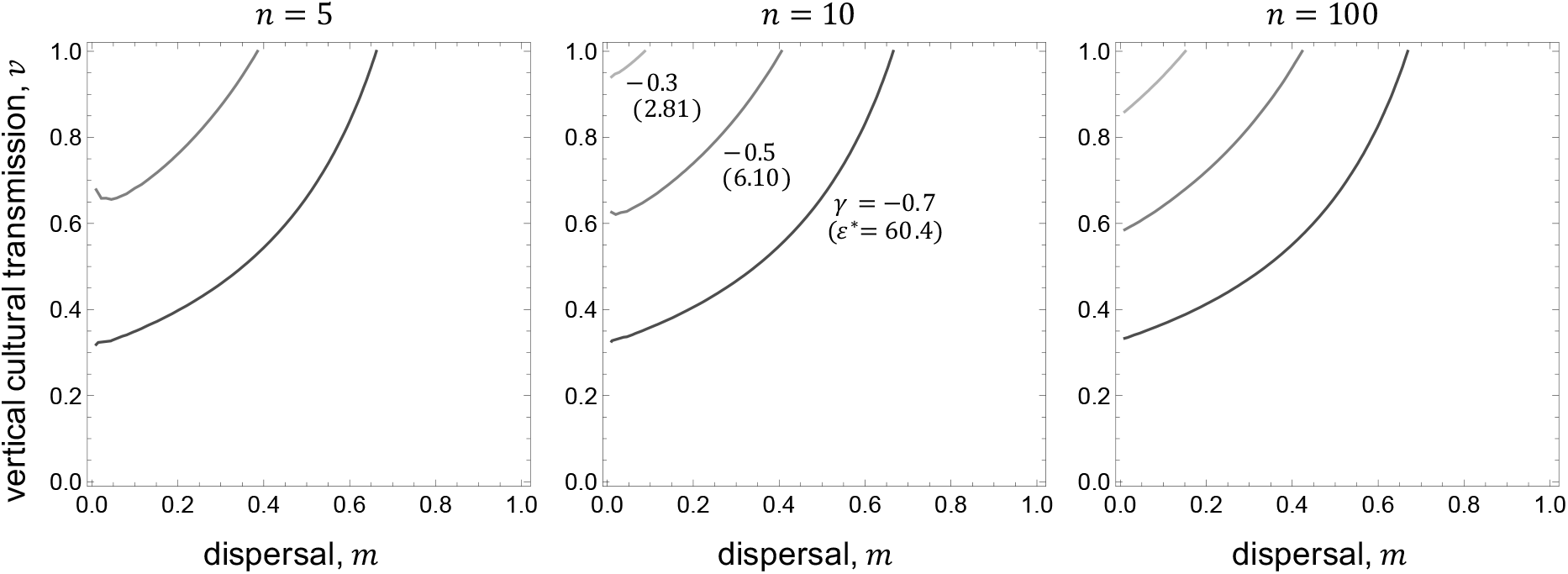
Levels of vertical cultural transmission and dispersal leading to the evolution of greatest knowledge. Each curve corresponds to the vertical cultural transmission, *υ*, and dispersal probability, *m*, under which selection favours *z*\* = *z*_MAX_ for a given level of synergy *γ* among social learners. For such *υ* and *m* values, gene-culture coevolution thus leads to the maximum level of knowledge 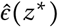 once the population expresses the evolutionary convergent strategy *z*\* (found by plotting the curve given by *s*(*z*_MAX_) = 0, with *z*_MAX_ defined by eq. (26) – parameter values: *β* = 2.5; *c* = *α* = 1; *n* = 5, 10, 100 from left to right panel; *γ* = −0.3, −0.5, −0.7 in increasingly dark gray; resulting 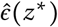 for each *γ* value shown on middle graph between brackets).

To check our analyses, we also performed individual based simulations that track gene-culture coevolution under the assumptions of our model (see Appendix E for details). We observed a very good match between these simulations and analytical predictions (Fig. 6), confirming our approach.

**Figure 6:**
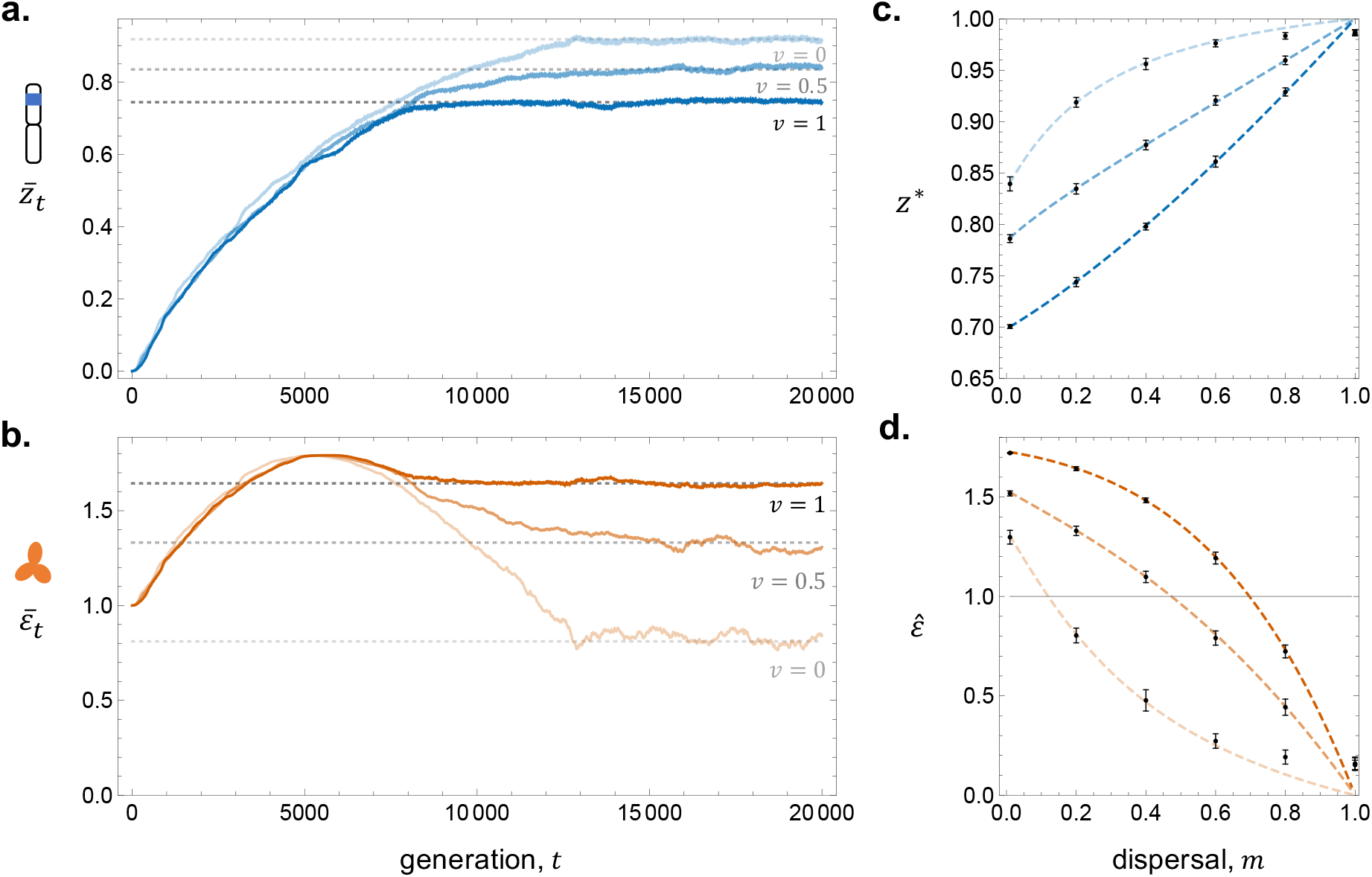
Individual based simulations versus analytical predictions of gene-culture coevolution. **(a,b)** Temporal dynamics of the population average investment into social learning ((a), blue) and concomitant population average knowledge ((b), orange) in simulated populations (full lines, see Appendix E for procedure), as well as predicted evolutionary convergent strategies ((a), dashed gray, computed by solving eq. (39) for *z*\* and equilibrium knowledge at evolutionary convergent learning strategy ((c), dashed gray, computed from eq. (25)), when *υ* = 0 (lighter shade), *υ* = 0.5 (intermediate shade), *υ* = 1 (darker shade);other parameters: *n* = 10, *m* = 0.2, *c* = 1, *α* = 1, *β* = 2.5, *γ* = 0, number of patches = 1000. **(c,d)** Predicted (dashed line) and simulated (black, population average calculated from generation 30000 to 50000 as points, error bars indicate standard deviation among generations) evolutionary convergent social learning strategies (c) and knowledge (d) as a function of dispersal (simulations with *m* = 0.01, 0.2, 0.4, 0.6, 0.8, 1) when *υ* = 0 (lighter shade), *υ* = 0.5 (intermediate shade), *υ* = 1 (darker shade); other parameters: same as (a,b).

## 5 Discussion

Through external modifications, individuals can readily impact the fitness of current but also future con-specifics (Dawkins, 1982; Lewontin, 1983; Odling-Smee et al., 2003; Bonduriansky, 2012; Govaert et al., 2019). As previously established, the selection gradient on a genetic trait with such extended effects depends on the effect of a trait change in one focal individual on the fitness of all current and future relatives in the population (Lehmann, 2007). So far, this kin selection approach has been applied to understand the joint evolutionary dynamics of traits together with the conditions they modify when populations are well-mixed (see below eq. (15) for connection and references), or when ecological inheritance is random within groups (see below eqs. (18)–(19)). In this paper we extended these results to allow ecological inheritance to be biased within groups due to non-random interactions, which is especially relevant to understand evolutionary driven changes in individual condition that can be preferentially transmitted from parent to offspring, such as micro-environments within patches (e.g., nests or burrows), or individuals characters (e.g., pathogen load, microbiome, culture).

Our analyses revealed how the combination of biased transmission of non-genetic traits and limited dispersal influence a variety of transgenerational pathways along which gene-driven modifications feed back on the fitness of relatives (eq. (10) onward). This variety reflects the multiple ways that can associate genetic variation to variation in non-genetic traits and fitness. A carrier of a genetic variant with extended effects may directly modify its own non-genetic trait, as well as indirectly those of current and downstream relatives living in its patch (eq. (12)). As a result, selection depends on trans-generational modifications cumulated over a genetic philopatric lineage (eqs. (13)–(15)) and how such modifications feed back on the fitness of members of this lineage (first term of eq. (10)). Due to social interactions within patches and oblique ecological inheritance, a carrier of a genetic variant can also modify the non-genetic trait of non-relatives within its patch even in the distant future. Selection then also depends on how this modification feeds back on the fitness of current and downstream relatives (second term of eq. (10), see also eqs. (16)–(19)).

As illustrated in our gene-culture coevolution example, our decomposition in terms of trans-generational kin selection effects allows to appreciate all the selective forces at play when individuals interact within patches, both directly and indirectly via extended effects. In addition, the selection gradient we have derived allows to investigate the quantitative effects of various genetic and ecological processes on the dynamics of traits coupled with the conditions they modify. In our example for instance, we found that provided some cultural information is passed vertically, populations with even moderate levels of dispersal (so that genetic relatedness within patches is low) can evolve a costly learning strategy that generates high levels of adaptive culture benefiting others (Figure 4, Supplementary Figure 1). Less intuitively perhaps, we further found that when social interactions have synergistic effects on social learning (equivalent to non-additive indirect genetic effects), evolution leads to greater levels of adaptive knowledge in the population under intermediate levels of both vertical transmission and dispersal (Figure 5). As these features are characteristic of human populations (van Schaik, 2016), it would be interesting to investigate in greater depth the nature of synergistic effects on learning, in particular their behavioural basis and evolution. More generally, this suggests that indirect extended genetic effects and ecological inheritance under limited gene flow interact in a non-trivial way, and that this interaction can have a significant impact on the evolution of extended traits.

Beyond gene-culture coevolution, our framework may offer insights into other evolutionary problems involving modifications of external features that can be transmitted between individuals. Of particular relevance are questions regarding host evolution to pathogens or microbiotic symbionts with mixed modes of transmission (Ebert, 2013). If much theory has highlighted the importance of kin selection emerging from spatial structure for such evolution, this theory typically focuses on random transmission patterns within spatial clusters (e.g., Brown and Hastings, 2003; Ferdy, 2009; Best et al., 2010; Horns and Hood, 2012; Débarre et al., 2012; Lion and Gandon, 2015). But preferential interactions between parents and their offspring are ubiquitous in social organisms, leading to vertically biased modes of transmission. Intuitively from our results, such transmission bias should favour the evolution of costly resistance to pathogens or maintenance of symbionts. This broad brush prediction aligns with population genetics models looking at the dynamics of host resistance (Schliekelman, 2007) or symbiotic alleles (Fitzpatrick, 2014). Unlike our mode however, these population genetics approaches do not allow for gradual transgenerational modifications to pathogens or symbionts owing to interactions with hosts.

Another relevant line of inquiry that can be pursued with our framework is the evolution of transmission bias itself. As the transmission function *F* in our model can depend on those individuals who pass their non-genetic trait (and those who inherit them, eqs. (1)–(2)), it is straightforward to investigate whether selection favours organisms to transmit (and/or receive) extended effects in a more vertical or oblique manner. Under gene-culture coevolution, models have studied how neutral evolution (Takahasi, 1999) and fluctuating environments (McElreath and Strimling, 2008) can lead organisms to rely on vertically-rather than obliquely-collected information when populations are well-mixed and information is fixed (i.e., does not accumulate). Here we suggest exploring how selection moulds the transmission of cumulative culture depending on limited dispersal. Selection in this case presumably depends on who controls the flow of adaptive information: parents should favour transmission to their offspring only, while offspring should favour whatever strategy maximising the information they receive. By homogenizing the genes and culture of individuals belonging to the same patch, limited dispersal can resolve this parent-offspring conflict and should therefore be pertinent for the evolution of transmission bias. Dispersal patterns and conflicts between interacting hosts should also be relevant to the evolution of host’s traits that influence the transmission of symbionts and pathogens. Existing literature on this topic has been mostly focused on the evolution of microbiotic strategies that favour one mode of transmission between hosts over another (e.g., Ferdy and Godelle, 2005; Boldin and Kisdi, 2012; Antonovics et al., 2017). Nonetheless hosts can also evolve remarkable strategies that influence transmission dynamics, such as altruistic suicide to limit kin exposure (Débarre et al., 2012; Berngruber et al., 2013; Humphreys and Ruxton, 2019). But the conditions that lead hosts to evolve behaviours favouring vertical over horizontal transmission or vice versa remain largely unexplored, especially for subdivided populations (Antonovics, 2017, for review). More generally, our framework may be useful to understand the coevolution of social behaviors with their evolutionary setting (Perc and Szolnoki, 2010; Akçay, 2020, for general remarks).

Although our model allows to better understand the nature of selection under a broad set of evolutionary scenarios involving extended genetic effects, it relies on several assumptions. Many of them, such as infinite population size, clonal reproduction of haploid genomes or rare mutation with weak effects are common to those of the adaptive dynamics framework and have been extensively discussed elsewhere (e.g., Geritz and Gyllenberg, 2005; Rueffler et al., 2006; Dercole and Rinaldi, 2008). One assumption that should be kept in mind is that transmission of non-genetic traits occurs after dispersal precludes strict maternal effects or more generally strict vertical ecological inheritance (as only philopatric individuals can be subject to these effects in our model). Put differently, extended genetic effects cannot disperse between patches in our model. In such a case, we expect kin effects on feedback selection to be weaker as individuals from different patches are less related than individuals from the same patch. In fact, a model of gene-culture coevolution showed that even moderate transmission of information between patches via dispersal can hinder the accumulation of adaptive culture (Ohtsuki et al., 2017). Extending our framework to include transmission before dispersal would therefore be an interesting avenue for future research. This would for instance offer the possibility to study host evolution in response to pathogens that can be directly transmitted from mothers to offspring in subdivided populations (Busenberg and Cooke, 1993, for epidemiological models in well-mixed populations), as well as dispersal evolution in response to pathogen load (Iritani and Iwasa, 2014). Another assumption we made that is limiting in the context of host-pathogen interactions is that patches have a fixed size. Our model therefore cannot track demographic changes due to infection. Such demographic changes are interesting from an epidemiological point of view but can also have pronounced effects on evolutionary dynamics (Lion and Gandon, 2015, for review). It would therefore be useful albeit challenging to include gene-driven demographic changes through extended effects to our conceptual model (Rousset and Ronce, 2004).

In sum, we have developed a theoretical framework to analyze how selection acts on genetic traits with extended effects that can be non-randomly transmitted across generations due to preferential interactions and limited dispersal. Our analysis disentangles the many paths via which selection can act due to limited dispersal and biased transmission, helping understand the nature of adaptation via trans-generational feedback effects between relatives. As illustrated by our gene-culture coevolution model, such feedback effects can affect evolutionary dynamics in significant and non-trivial ways. More broadly, our theory can help us understand the interplay between genetic and extra-genetic ecological inheritance, with implications for how organisms evolve to transform their culture, microbiome and external environments.

## Acknowledgements

This research was conducted thanks to a visiting grant to SOKENDAI to CM and HO. CM is supported by a SNSF grant (No. PCEFP3181243). JYW and HO are supported by JSPS KAKENHI (No.16K07524). JYWis supported by JSPS KAKENHI (No.16K05283 and 16H06412). HO is supported by JSPS KAKENHI (No.16H06324).

## Appendices

## A Basic reproductive number in the island model of dispersal

In this appendix, we specify the reproductive number *R*_0_(*z*^m^, *z*), which lies at the basis of our analysis. In the island model of dispersal, the reproductive number *R*_0_(*z*^m^, *z*) (sometimes referred to as “lineage fitness proxy”) is defined as the expected number of successful offspring produced by an individual that is randomly sampled from a local lineage of rare mutants with genetic trait *z*^m^ in a resident population with genetic trait *z* (see Mullon et al., 2016, for homogeneous groups; and Lehmann et al., 2016, for heterogeneous groups).

To connect the above definition of *R*_0_(*z*^m^, *z*) with our model, let us first introduce *M_t_* ∈ {0,1,2,…, *n*} as the random variable for the number of mutant individuals with trait *z*^m^ at generation *t* = 0, 1, 2, … in a mutant patch (i.e., a patch in which the mutant arose as a single copy at time *t* = 0). As the mutant arose as a single copy at time *t* = 0, we have *M*_0_ = 1. The remaining *n* – *M_t_* individuals in this mutant patch at generation *t* are residents with genetic trait *z*. However, these resident individuals can express different extended traits than residents from resident patches due to their interactions with genetic mutants (via indirect extended genetic effects and oblique ecological inheritance). To capture this variation among carriers of the resident genetic trait in the mutant patch, we distinguish between different cohorts of residents, varying in when their ancestors first arrived in the mutant patch. Specifically, we let *R*_*t*,0_ denote the number of residents who had an ancestor in the mutant patch when the first mutant arose at time *t* = 0, and *R_t, t−h_* be a random variable for the number of residents at generation *t* in the mutant patch whose local lineage was initiated exactly *h*(< *t*) generations ago. By definition, these random variables sum to the total number of residents at time *t*, 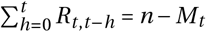 (and as there is a single mutant at time *t* = 0, *R*_0,0_ = *n* − 1).

With the above notation, the relevant genetic-demographic state *S_t_* of the mutant patch at time *t* is described by a non-negative partition of the integer *n* into *t* + 2 distinctive cohorts:

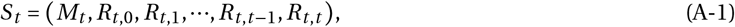

where 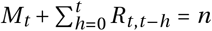. A realization of the stochastic process from time 0 to *t* is thus characterized by a collection of such partitions, which we denote by 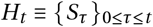, i.e., *H_t_* describes the genetic-demographic history of the mutant patch from time 0 to *t*. We let 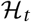 represent the set of all possible genetic-demographic histories from time 0 to *t* in a mutant patch (so that 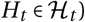), and Pr(*H_t_*) be the probability that the history *H_t_* is realized.

The probability that an individual randomly sampled from the mutant lineage (over the lifetime of this lineage in the patch) is sampled at time *t* with patch history *H_t_* is then given by

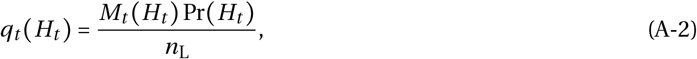

where we have deliberately stressed the trivial dependence of *M_t_*(*H_t_*) on *H_t_*, and

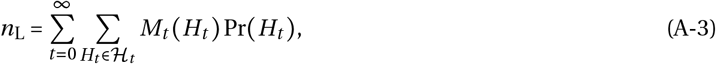

is the expected cumulative size of the local mutant lineage over its lifetime (note that owing to the constant influx of resident immigrants into a mutant patch, extinction of a local mutant lineage is certain, i.e., lim_*t*→∞_ Pr(*M_t_* = 0) = 1, so that *n*_L_ < ∞ is bounded;see eq. A19 in Mullon et al., 2016, for equivalence with eq. (A-2) in the absence of historical effects). For our analysis, it will be sometimes convenient to re-write eq. (A-2) as

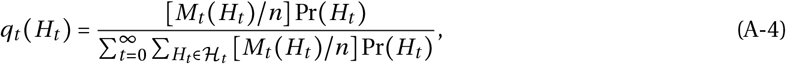

which is proportional to the fraction *M_t_*(*H_t_*)/*n* of mutants in the mutant patch at time *t*.

Let us now consider the fitness of such a randomly sampled mutant (so at time *t* with patch history *H_t_*). Using the fitness function eq. (6) from the main text, the fitness of this focal mutant can be written as

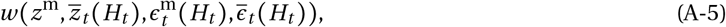

where *z*^m^ is the genetic trait value of the mutant;

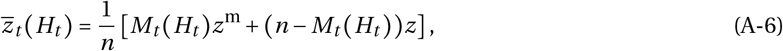

is the average trait value in its patch; 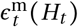 denotes the extended trait of the focal mutant and 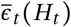 the average of extended trait in the patch, both of which depend on the genetic-demographic history of the mutant patch (we will specify these dependencies later in Appendix C).

From the above considerations, we can write the expected fitness of a representative mutant (randomly sampled from a local lineage of rare mutants), i.e., the basic reproductive number as,

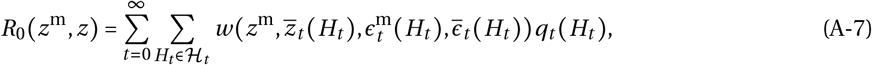

or equivalently as

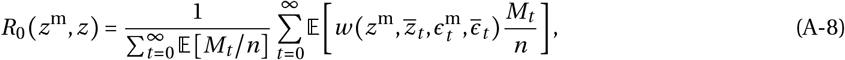

where expectation is taken over the probability distribution Pr(*H_t_*) (to avoid notational burden, we occasionally drop the dependency on *H_t_* of any function *f*(*H_t_*) and simply write it *f* = *f*(*H_t_*), as in eq. (A-8)).

## B Deriving the selection gradient

In this appendix, we derive the selection gradient eqs. (8)–(10) of the main text.

### B.1 Decomposing the selection gradient

By plugging the basic reproductive number eq. (A-7) into the selection gradient eq. (7), we obtain

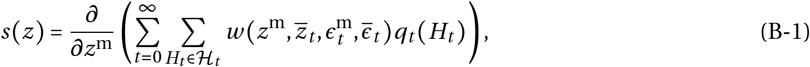

where here and thereafter, all derivatives are estimated at the resident genetic trait *z* and equilibrium extended trait 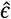. Using the chain rule, the selection gradient can then be expressed as

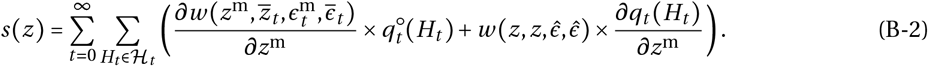

Here, 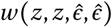 is individual fitness in a monomorphic population of residents, and 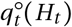 is the probability that under neutrality (so when mutants and resident have the same genetic trait value, *z*^m^ = *z*), a randomly sampled individual from a local mutant lineage is sampled at time *t* with patch history *H_t_*, i.e.,

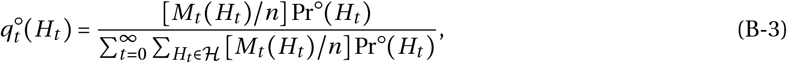

where Pr°(*H_t_*) is the probability that the history *H_t_* is realized under neutrality.

We can then use the fact that fitness in a monomorphic population is one as the total population size is constant (i.e., 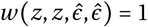), and that *q* satisfies 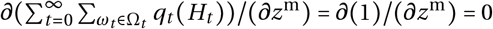 (see eq. (A-2)–(A-3)). Therefore the second term in eq. (B-2) vanishes and we obtain

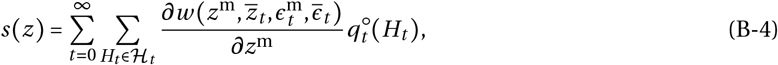

or equivalently,

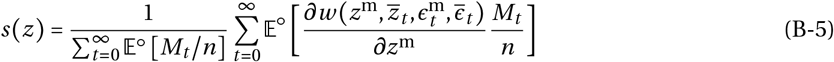

where 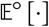 represents expectation taken over Pr°(*H_t_*).

We then expand the fitness derivative that appears in the selection gradient eq. (B-4) using the chain rule as

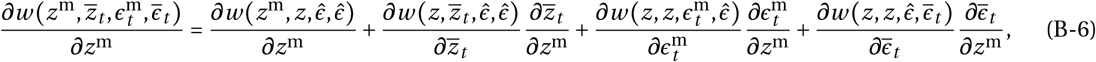

where the first two summands capture the genetic effects on fitness and the other two, the feedback effects through the extended trait expressed by the focal individual whose fitness is being considered and through the average non-genetic trait expressed in its patch.

Substituting eq. (B-6) into eq. (B-4) allows us to decompose the selection gradient as the sum of two terms (as in the main text eq. (8)),

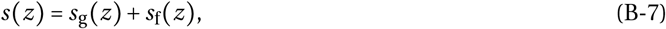

where

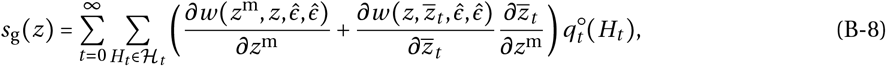

captures selection on *z* according to its effects on fitness, and

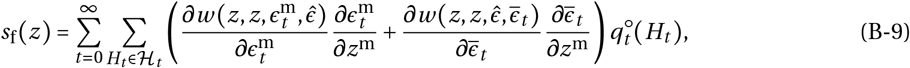

according to the way it feedbacks on fitness via the non genetic trait *ϵ*. We specify both of these selection terms in the next two sections.

### B.2 Genetic effects on fitness

From the definition of the average genetic trait in the patch 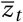 (eq. A-6), we have 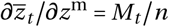. Substituting for this into eq. (B-8), we find that selection on *z* according to its effects on fitness can be re-written as

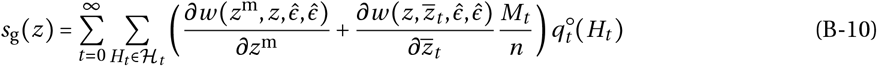

As the fitness derivatives are independent of time when they are evaluated at resident *z* and 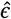, they can be taken out of the expectation in eq. (B-10), yielding

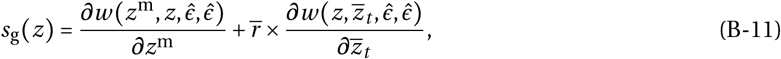

where

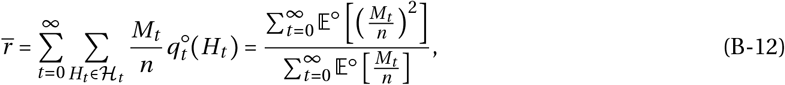

is the probability that under neutrality, a random individual sampled from the patch of a representative mutant (i.e., a randomly sampled from a local lineage of rare mutants) is also a mutant (from the definition of 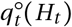, eq. (A-2)). Since there is no distinction between mutants and residents under neutrality (*z*^m^ = *z*), the quantity 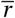 defined by eq. (B-12) is equal to the probability that two individuals sampled with replacement from the same patch at the same generation belong to the same local lineage, i.e., the probability that they carry an allele that is identical-by-descent at a neutral locus, which corresponds to the definition of patch-average relatedness (e.g., Rousset, 2004). Thus, we find as required that eq. (B-11) is eq. (9) of the main text.

### B.3 Feedback effects

We now turn our attention to feedback selection effects, eq. (B-9), which can be re-written as eq. (10) of the main text,

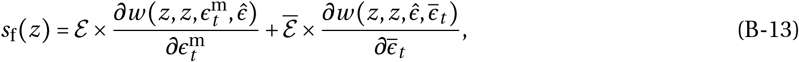

where

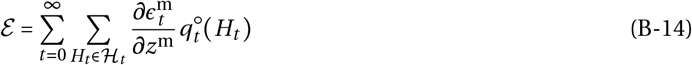

is by definition the influence of the genetic mutant on the expected extended trait expressed by a representative member of the local mutant lineage (i.e., randomly sampled from the local mutant lineage), and similarly,

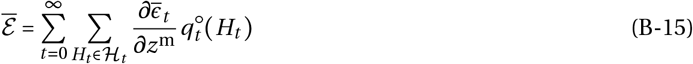

is the influence of the genetic mutant on the expected average extended trait in the mutant patch experienced by a representative mutant. We evaluate these expectations in Appendix C.

## C Effects of a genetic mutant on extended traits

Here, we evaluate the effects of a genetic mutant on extended traits that are under selection, 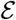 and 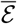 (eqs. (B-13)–(B-15)) and derive eqs. (11)–(19) of the main text. To do so, we first characterise the dynamics of extended traits in a mutant patch under the assumptions laid out in section 2.2 of the main text.

### C.1 Dynamics of extended traits

#### C.1.1 Mutant extended trait

As the genetic mutant is globally rare, any carrier of this mutation in the mutant patch is necessarily philopatric in the island model of dispersal. Using eq. (1) of the main text, the extended trait expressed by a carrier of the genetic mutation at generation *t* in the mutant patch is thus given by

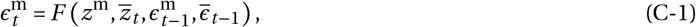

where recall that *z*^m^ is the genetic trait carried by the focal mutant; 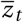, the average genetic trait in the mutant patch at generation *t* (see eq. A-6); 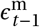, the extended trait expressed by the parent of this mutant at generation *t*−1 (which is necessarily in the same patch as we are considering a local mutant lineage);and 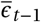, the average extended phenotype at the previous generation (which is characterised in eq. (C-4)).

#### C.1.2 Resident extended trait in the mutant patch

The extended trait of individuals carrying the resident genetic trait *z* can be modified by the genetic mutant *z*^m^ via indirect extended genetic effects and oblique ecological inheritance. We therefore also need to characterize the mutant-induced dynamics of the extended traits of carriers of the resident trait *z* in the mutant patch. These dynamics are complicated as, in contrast to carriers of the genetic mutant, individuals carrying the resident genetic trait *z* in the mutant patch may be philopatric or immigrants. In addition, effects of the genetic mutant on the extended trait of a resident individual will depend on when the local lineage of this resident individual was initiated into the mutant patch (i.e., when the ancestor of a resident individual first immigrated in the mutant patch). We thus have to consider different classes of resident individuals in the mutant patch.

##### New resident immigrant

Let us first consider a carrier of the resident trait *z* at generation *t* that has just immigrated into the focal patch. By 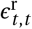, we denote the extended trait of a carrier of the resident at generation *t* (first subscript of 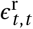) that has immigrated into the mutant patch at generation *t* (second subscript of 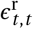; as a reminder, the number of such carriers is *R_t,t_*, see Appendix A). From eq. (2) of the main text, the extended trait of such a new resident immigrant is,

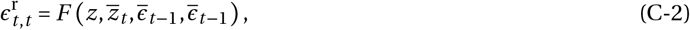

which reflects that a new immigrant cannot experience vertical ecological inheritance (as its genetic parent is absent from the patch it lands in).

##### Philopatric resident

By contrast, philopatric carriers of the resident genetic trait do have a local parent and their extended trait can therefore be preferentially transmitted vertically from their parent. However, the dynamics of the extended trait along such a local resident lineage depends on precisely when this lineage was initiated, i.e., when its first local ancestor immigrated into the mutant patch. To distinguish between these different cohorts of residents, we let 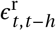 denote the extended trait of a carrier of the genetic resident at generation *t* whose local lineage in the mutant patch was initiated 1 ≤ *h* ≤ *t* generations before (as a reminder, the number of such carriers is *R_t,t−h_*, see Appendix A). Using eq. (1) ofthe main text, such extended trait is given by

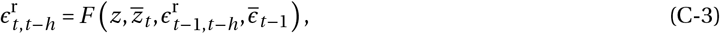

so that at any generation, carriers of the resident genetic trait can potentially express many different extended traits in the mutant patch, depending on the history of the interactions between their local lineage and the lineage of the mutant.

#### C.1.3 Average extended trait in the mutant patch

For our analysis of selection, we further need to characterise, 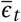, the average extended trait in the mutant patch at time *t*. From the above definitions (eqs. (C-1)–(C-3)), the patch average extended trait is given by

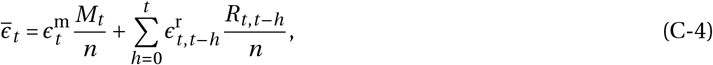

where recall that 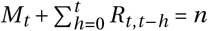.

### C.2 Conditional effects of the genetic mutant on extended traits in the mutant patch

Next, we use eqs. (C-1)–(C-4) to characterize the effect of the genetic mutant *z*^m^ on the genetic traits expressed in the mutant patch at an arbitrary generation *t*, conditional on a specific sequence of genetic-demographic states *H_t_*. Specifically, our goal here is to characterize:

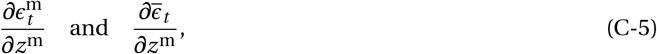

given a realized sequence *H_t_* in the mutant patch. We will then marginalise these effects over the relevant mutant-experienced distribution, 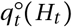, of such genetic-demographic states *H_t_* to finally obtain 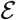 and 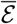 in section C.3 (eqs. (B-14)–(B-15)).

First, we take the derivative of both sides of equations (C-1), (C-2) and (C-3) to obtain the following recursions for the effect of the genetic mutant on extended traits in the mutant patch,

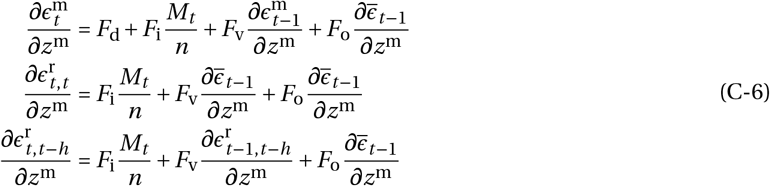

for 1 ≤ *h* ≤ *t* (with *F*_d_, *F*_i_, *F*_v_ and *F*_o_ defined in eq. (4) and eq. (5);all these effects depend on resident *z*, but we do not write these dependencies explicitly in the appendix). Similarly, by taking the derivative of both sides of eq. (C-4), the effect on the average extended trait at time *t* is

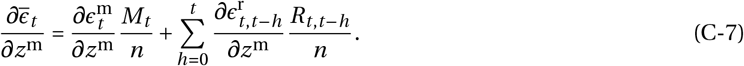

The dynamical system described by eq. (C-6) has initial conditions,

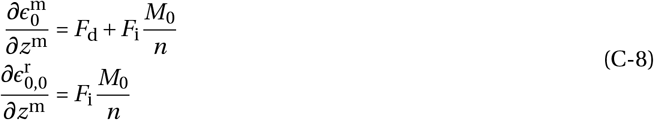

where *M*_0_ = 1 as a single mutant appears in the mutant patch at time *t* = 0. Combining eqs. (C-7)–(C-8), the initial condition of the average effect is

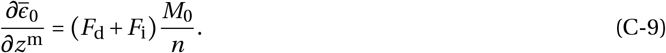

Given a realized sequence of genetic-demographic states *H_t_*, eqs. (C-6)–(C-9) describe a discrete dynamical system in time *t* whose solution gives us eq. (C-5). In principle, one could therefore solve this system and then marginalise the solution over 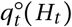 to obtain 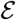 and 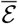 (eqs. (B-14)–(B-15)). However, given the dimensionality of the dynamical system we are considering (up to *t*+2 dimensions at each time step *t* = 0,1,…), such a strategy seems to be extremely complicated. We will therefore rely on an alternative argument, as elaborated in the next section.

### C.3 Unconditional effects of the genetic mutant on extended traits in the mutant patch

We proceed to marginalise eq. (C-5) over 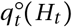 using eqs. (C-6)–(C-9) to obtain 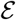 and 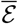 (eqs. (B-14)–(B-15)). To do so, note first that using eq. (B-3), we can re-write 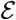 and 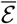 as

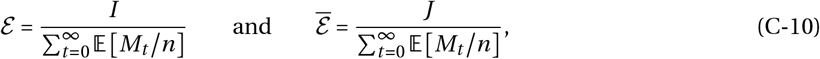

where 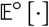 represents expectation taken under neutrality (over Pr°(*H_t_*)), and

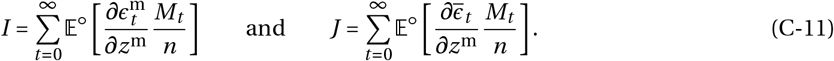

Our strategy is to first derive *I* and *J*, which we will then plug back into eq. (C-10).

We start by characterising *I*. Multiplying both sides of the equation on the first line of eq. (C-6) by *M_t_*/*n* and taking expectation under neutrality, we obtain

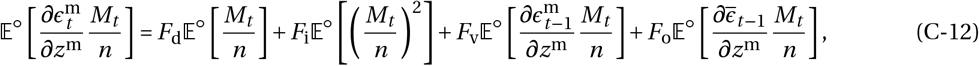

for *t* ≥ 1. Let us first focus on the last two summands of the above equation. Both consist of the expected value of a product of two random variables: one that depends on events up until time *t* − 1, 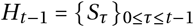 (the random variables 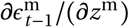 and 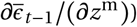); and one on events at time *t*, *S_t_* (the random variable *M_t_*/*n*).

We can therefore disentangle these using general properties of conditional expectation,

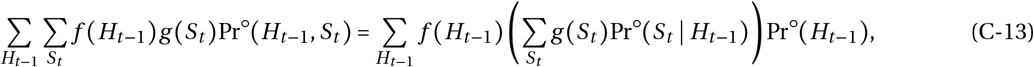

where *f*(*H_t−1_*) corresponds to 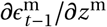 and 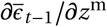, and *g*(*S_t_*) corresponds to *M_t_/n* in eq. (C-12). Equivalently, we have

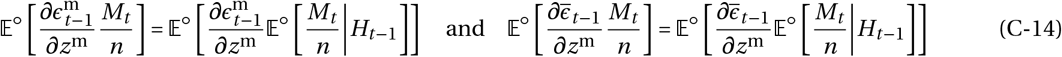

for the last two summands of eq. (C-12).

Next, note that under neutrality, the conditional probability distribution, Pr°(*S_t_* | *H_t−1_*), of *S_t_* = (*M_t_*, *R_t,0_*,⋯, *R_t,t_*) given the genetic-demographic history *H_t−1_* is a multinomial distribution,

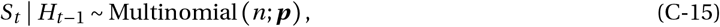

where ***p*** is a (*t* + 2)-dimensional vector of probabilities given by

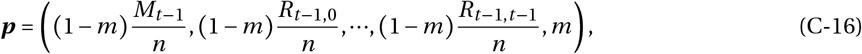

with *m* as the backward probability of dispersal, i.e., the probability that under neutrality, an individual is an immigrant. Using the formula for the expectation of a multinomial distribution, we then have

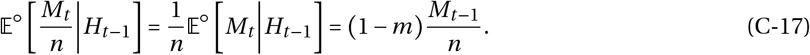

Substituting eq. (C-17) into eq. (C-14) which is in turn plugged into eq. (C-12), we obtain

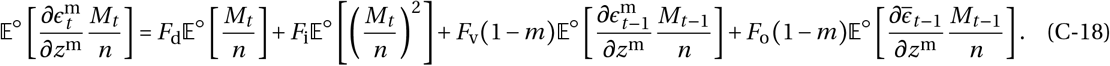

Summing the above equation from *t* = 1 to infinity and using eq. (C-11), we get

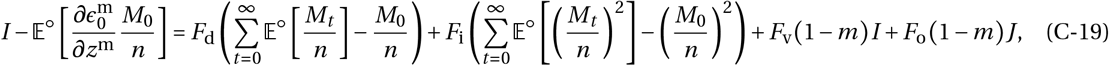

but note that from eq. (C-8),

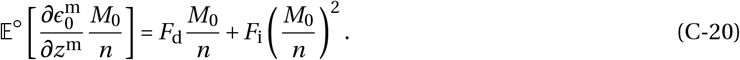

Substituting this into the lefthand side of eq. (C-19), we obtain

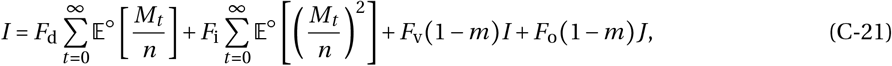

after some re-arrangements. Then, note that by dividing both sides of eq. (C-21) by 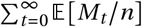, we get an identity in terms of 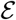 and 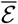,

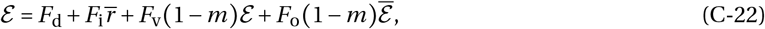

using eqs. (B-12) and (C-10).

We proceed similarly to derive an identity for 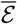 from *J* (eq. (C-11)). First, we multiply both sides of eq. (C-7) by *M_t_/n* to give,

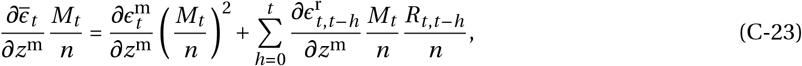

into which we substitute eq. (C-6) to obtain,

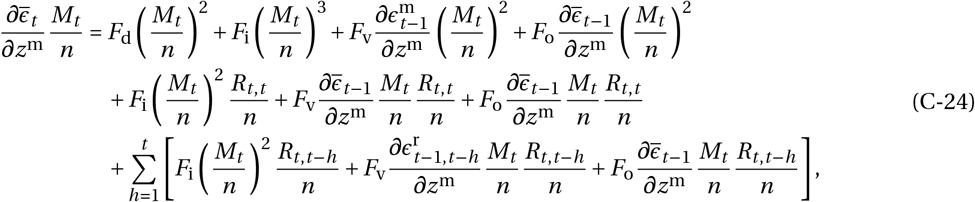

for *t* ≥ 1. Re-arranging the above and using the fact that 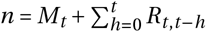, we get,

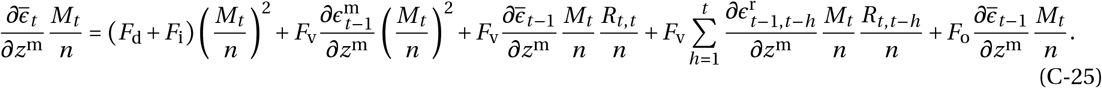

Taking the expectation of both sides of eq. (C-25) leads us to

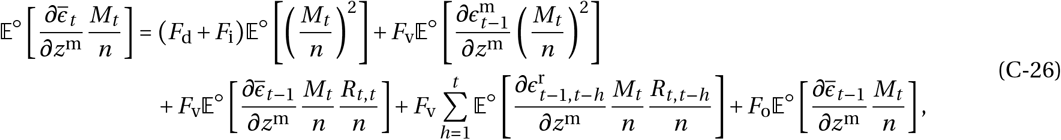

which can be rewritten as

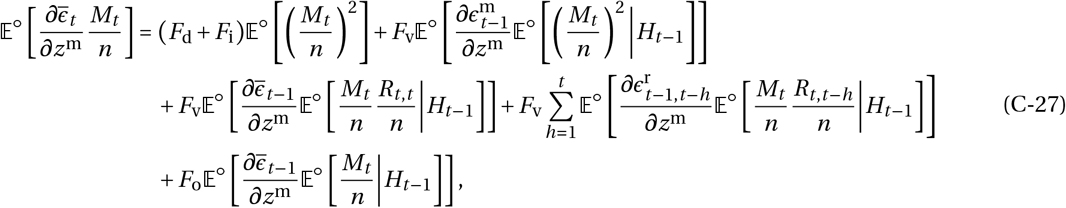

using the properties of conditional expectation (eq. (C-13)).

We can then use eq. (C-15) to evaluate the conditional expectations that appear in eq. (C-27),

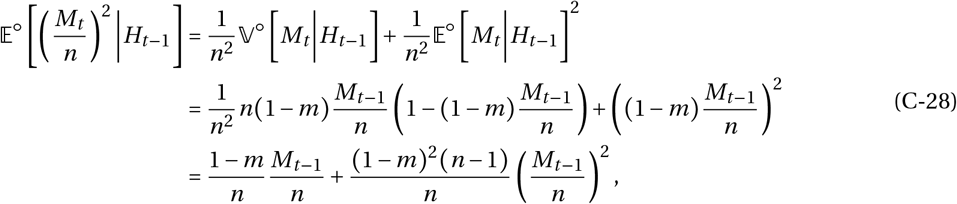

where 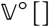 represents variance with respect to 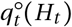. Similarly, using the formula for the covariance between random variables that have a multinomial distribution, we have

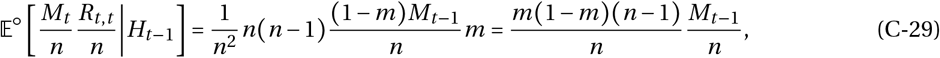

and

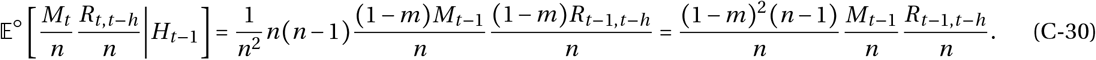

Substituting eqs. (C-17) and (C-28)–(C-30) into eq. (C-27) and some re-arrangements allows us to write,

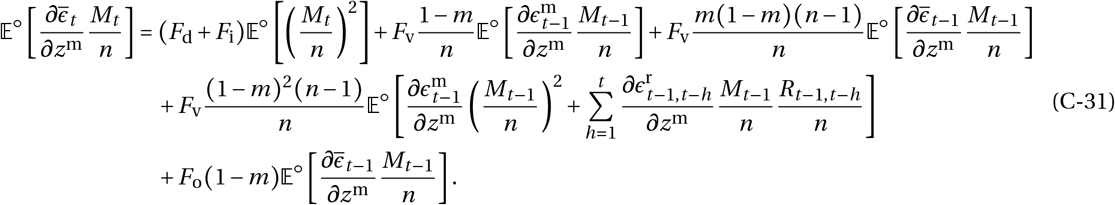

However, note that from eq. (C-7), the term within the expectation operator on the second line of eq. (C-31) can be re-written as

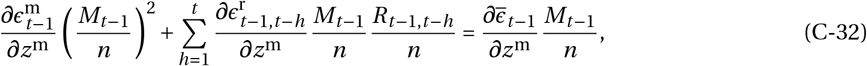

yielding

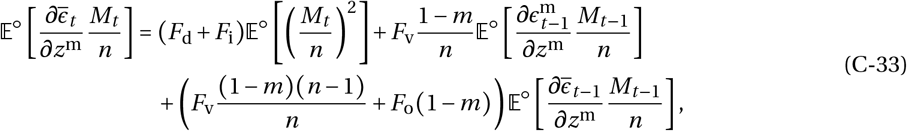

for eq. (C-31). Summing both sides of eq. (C-33) from *t* = 1 to infinity and using eq. (C-11) then gives us

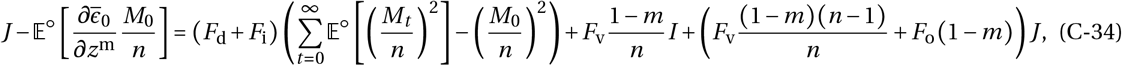

which using the fact that

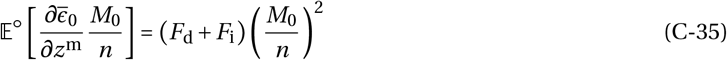

(from eq. (C-9)) becomes

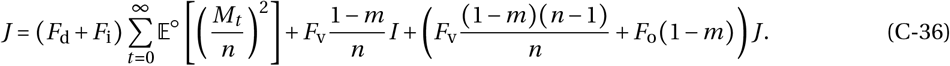

Finally, by dividing both sides of eq. (C-36) by 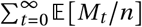 and using eqs. (C-10) and (B-12), we get,

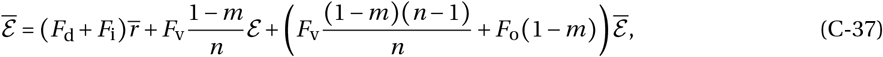

i.e., an identity in terms of 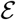 and 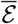.

Solving eqs. (C-22) and (C-37) simultaneously for 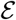 and 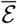, we eventually get,

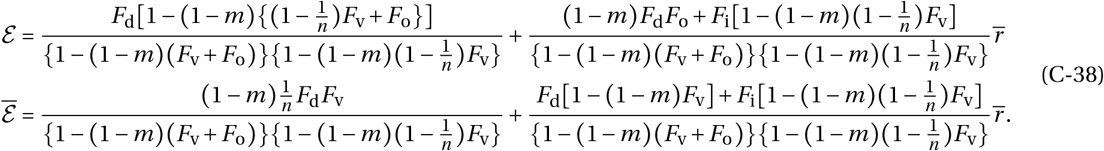

which are equivalent to eqs. (11)–(14) and (16)–(19) of the main text, in which we have decomposed eq. (C-38) into different components according to effects that occur within- and between-generations, as well as effects due to deterministic or stochastic genetic fluctuations. To see how we obtain this decomposition, consider first eqs. (C-22) and (C-37) in the absence of trans-generational effects (i.e., no ecological inheritance, *F*_v_ = *F*_o_ = 0),

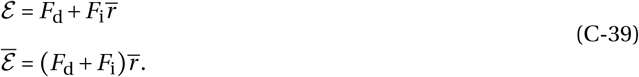

These are precisely the intra-generational effects, 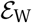 and 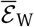, given in the main text (eq. (12) and (17)). Second, let us ignore stochasticity stemming from genetic fluctuations in the mutant patch, i.e., instead of eqs. (C-28)–(C-30), we use a deterministic approximation,

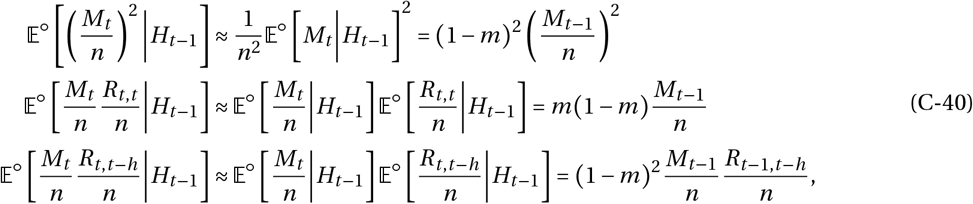

so that in effect, we ignore variance in genetic-demographic state in the mutant patch and just consider deterministic changes in state. Substituting eqs. (C-17) and (C-40) into eq. (C-27) and following the same argument as above, we eventually obtain the system

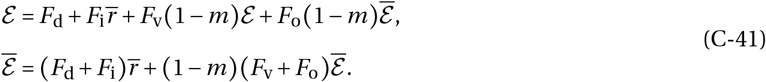

The solution of this system is

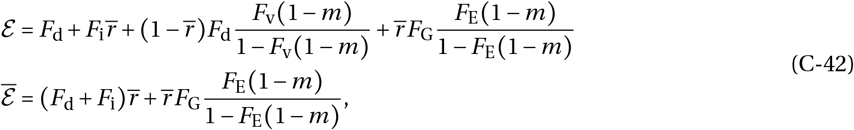

i.e., the sum of within-generation effects (eq. (C-39)) and deterministic trans-generational effects as described in the main text (eq. (13) without Δ_*ϵ*_(*z*) and eq. (18) without 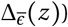). Finally, deducting eq. (C-42) from eq. (C-38), we are left with Δ_*ϵ*_(*z*) and 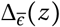 as given in the main text, eqs. (14) and (19), which thus capture trans-generational effects due to stochastic fluctuations in the genetic-demographic state of the mutant patch.

## D Trans-generational feedback effects of social learning

The functions, *ϕ*(*z*) and *ψ*(*z*), which feature in the expression for the trans-generational effects of social learning eq. (38) of the main text, are given by

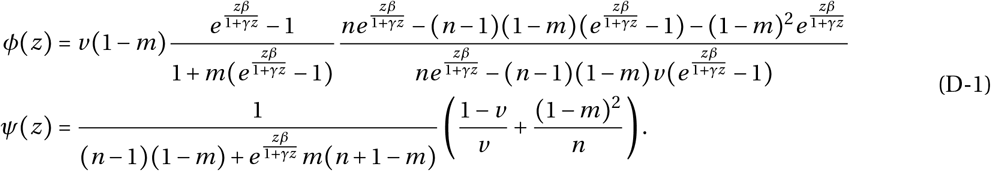

Showing that *ϕ*(*z*) ≥ 0 and *ψ*(*z*) ≥ 0 can be easily done with an algebraic computer program (e.g., using the function Reduce[] in Wolfram Research, Inc., 2016).

## E Individual based simulations

We performed individual based simulations for a population composed of *n*_d_ = 1000 groups, each populated by *n* = 10 individuals, using Mathematica 11.0.1.0 (Wolfram Research, Inc., 2016). Our simulation code can be downloaded from the public repository XXX (determined upon acceptance for publication). Each adult individual *i* ∈ {1,…, *n*_d_ *n*} is characterized by its genetic trait *z_i_* for social learning and its level of knowledge *ϵ_i_*. Starting with a monomorphic population with *z_i_* = 0 (i.e., no social learning) and *ϵ_t_* = 1 for all *i*, we track for 50000 generations the evolution of the phenotypic distribution for social learning as well as the distribution of knowledge level under the influx of genetic mutations. Each generation is composed of three steps:

i. *Reproduction*. At the beginning of a generation, we first calculate the fecundity *f_i_* of each individual *i* according to its trait *z_i_* and knowledge *ϵ_i_* (using eq. 27, see Fig. 6 for parameter values). Then, we form the next generation of adults by sampling *n* individuals in each group with replacement from the whole pool of parents according to parental fecundity (multinomial sampling), but to capture limited dispersal, the fecundity of each individual from the parental generation is weighted according to whether or not they belong to the group on which the breeding spot is filled: if an individual belongs to the same group in which a breeding spot is filled, its weighted fecundity is *f_i_*(1 − *m*), where *m* is the dispersal probability; if it belongs to another group, its weighted fecundity is *f_i_m*/(*n*_d_ − 1) (as a disperser is equally likely to reach any other group, it lands with probability 1/(*n*_d_ – 1) in a focal group).
ii. *Genetic mutation*. Once an individual is chosen to fill the breeding spot, its genetic trait mutates with probability 0.01, in which case we add to parental genetic value a perturbation that is sampled from a normal distribution with mean 0 and variance 0.005. The resulting phenotypic value is truncated to remain between 0 and 1.
iii. *Learning*. The final step within a generation *t* consists of updating the knowledge *ϵ_i_* of new individuals. If an individual *i* with genetic trait *z_i_* is philopatric, its knowledge is 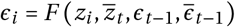 where *F* is given by eq. (22) (see Fig. 6 for parameter values), with 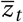 as the average genetic trait among individuals in its group, *ϵ_t−1_* is the knowledge of the parent of individual *i*, and 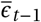 is the group-average knowledge in the parental generation of its group. If however individual *i* is an immigrant, its knowledge is 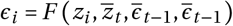, where 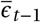 is the the group-average knowledge in the parental generation of the group in which individual *i* settles. This completes one generation, after which we return to step (i).

## Supplementary Figures

**Figure 1:**
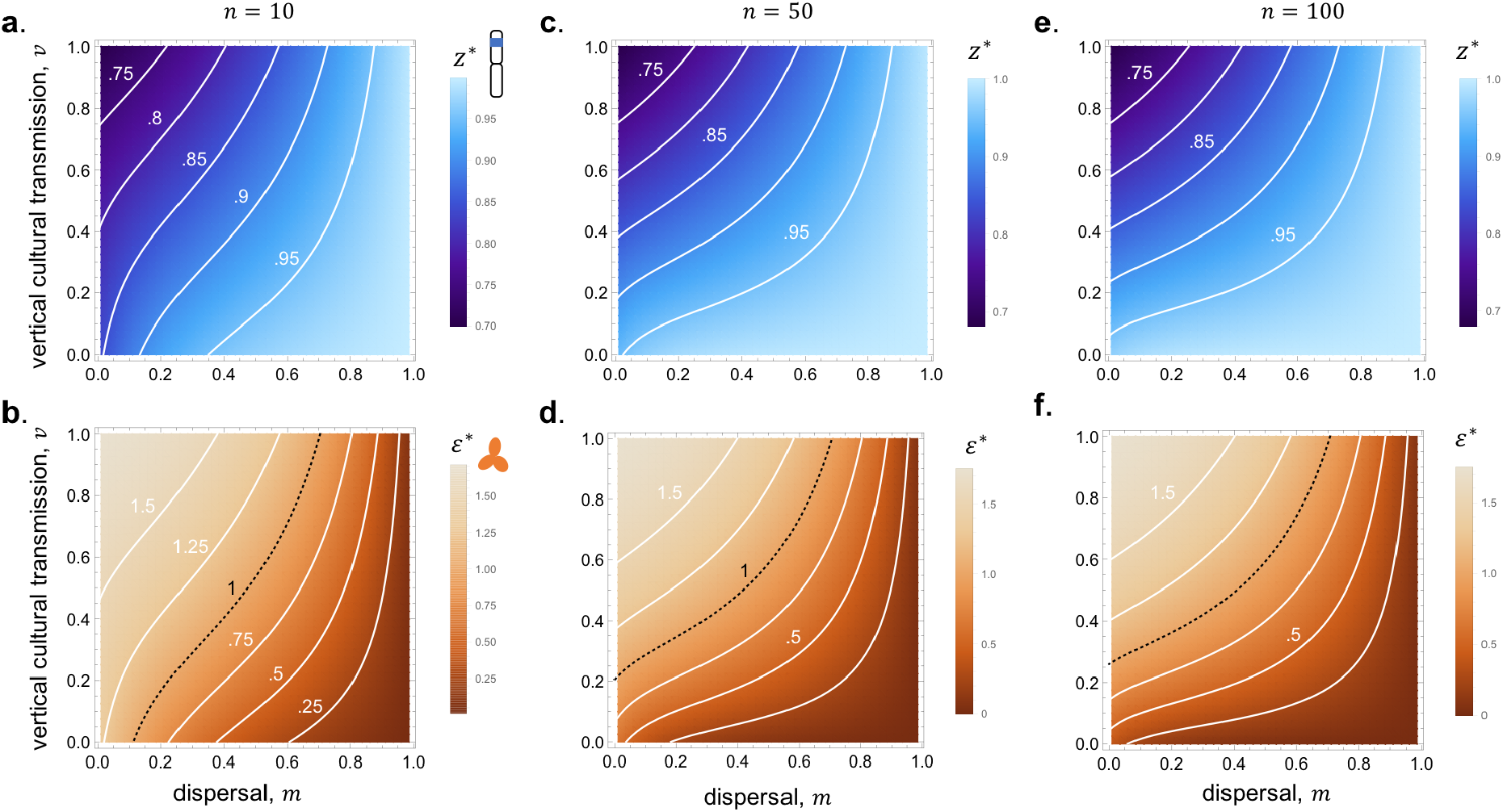
Effect of group size on evolutionary convergent learning strategy and associated adaptive knowledge. **Top row (a,c,e):** Contours of evolutionary convergent learning strategy *z*\* according to vertical cultural transmission on y-axis and limited gene flow on x-axis (computed by solving numerically eq. (39) for *z*\*, parameter values: *γ* = 0, *β* = 2.5; *c* = *α* = 1;(a) *n* = 10; (c) *n* = 50; (e) *n* = 100;lighter colours mean greater investment *z*\* in social learning – so less individual learning – see colour figure legend). **Bottom row (b,d,f)**: Contours of equilibrium knowledge 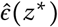 at evolutionary convergent learning strategy *z*\* according to vertical cultural transmission on y-axis and limited gene flow on x-axis (computed from values found in top row for *z*\* and eq. (25), same parameters as top row). Contour for 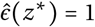, which is the threshold above which cumulative culture occurs (see Fig. 2c), shown as a black dashed line (lighter colours mean greater knowledge in the population, see colour figure legend).

## Footnotes

* Because the genetic average 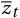 includes the focal genetic trait *z*_•_ and because the extended average 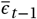 includes the extended trait of its parent *ϵ*_*t*−1_, *F* does not completely distinguish between direct and indirect genetic effects or vertical and oblique ecological inheritance; such formulation simplifies mathematical analysis and all our results can be straightforwardly applied to cases where ecological inheritance is strictly oblique or extended genetic effects are strictly indirect by correctly defining the relevant averages in term of focal phenotype and that of its neighbours, see eq. (20) for e.g.

† We can express the average fecundity in the patch as a function of average traits 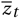 and 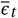 as variation in these is assumed to be small.

## Notes

### Competing Interest Statement

The authors have declared no competing interest.

